# Global parameter optimisation and sensitivity analysis of antivenom pharmacokinetics and pharmacodynamics

**DOI:** 10.1101/2023.03.13.532354

**Authors:** Natalie M Morris, Johanna A Blee, Sabine Hauert

## Abstract

In recent years it has become possible to design snakebite antivenoms with diverse pharmacokinetic properties. Owing to the pharmacokinetic variability of venoms, the choice of antivenom scaffold may influence a treatment’s neutralisation coverage. Computation offers a useful medium through which to assess the pharmacokinetics and pharmacodynamics of envenomation-treatment systems, as antivenoms with identical neutralising capacities can be simulated. In this study, we simulate envenomation and treatment with a variety of antivenoms, to define the properties of effective antivenoms. Systemic envenomation and treatment were described using a two-compartment pharmacokinetic model. Treatment of *Naja sumatrana* and *Cryptelytrops purpureomaculatus* envenomation was simulated with a set of 200,000 theoretical antivenoms across 10 treatment time delays. These two venoms are well-characterised and have differing pharmacokinetic properties. The theoretical antivenom set varied across molecular weight, dose, k_on_, k_off_, and valency. The best and worst treatments were identified using an area under the curve metric, and a global sensitivity analysis was performed to quantify the influence of the input parameters on treatment outcome. The simulations show that scaffolds of diverse molecular formats can be effective. Molecular weight and valency have a negligible direct impact on treatment outcome, however low molecular weight scaffolds offer more flexibility across the other design parameters, particularly when treatment is delayed. The simulations show k_on_ to primarily mediate treatment efficacy, with rates above 10^5^ M^-1^s^-1^ required for the most effective treatments. k_off_ has the greatest impact on the performance of less effective scaffolds. While the same scaffold preferences for improved treatment are seen for both model snakes, the parameter bounds for *C. purpureomaculatus* envenomation are more constrained. This paper establishes a computational framework for the optimisation of antivenom design.

## 1 Introduction

Snakebite envenomation is a neglected tropical disease which is responsible for approximately 81,000 – 138,000 deaths and 400,000 cases of disability each year (Gutiérrez et al., 2017; World Health Organisation, 2021). The vast majority of snakebite morbidity and mortality occurs in rural developing regions, leading to a significant socioeconomic and health burden in these areas. Timely access to specific, safe, and effective antivenoms would however prevent death and many of the other ill-effects of snakebite (Gutiérrez et al., 2017).

The diverse symptoms of envenomation are produced by the protein toxins present within venom. The toxin profiles of venoms from different snake species can vary substantially (Tasoulis and Isbister, 2017). Venom toxins may act locally and/or systemically, are often highly synergistic, and can disrupt targets within the blood and vasculature, the nervous system, soft tissue, organs, and more (Ferraz et al., 2019; Gutiérrez et al., 2017; Laustsen, 2016). Venom toxins can vary considerably in size, ranging from small peptides to large multi-subunit enzymes (Ferraz et al., 2019; Tasoulis and Isbister, 2017). Proteins of different sizes, shapes, and isoelectric points have different routes and rates of absorption, elimination, and biodistribution throughout the body (Bumbaca et al., 2012; Deen et al., 1979; Sánchez-Félix et al., 2020). This imparts pharmacokinetic variability to the toxins within different venoms (Gutiérrez et al., 2003; Sanhajariya et al., 2018). It has therefore been suggested that envenomation treatment efficacy may be improved through the use of antivenoms with similar pharmacokinetic properties to their targeted toxins (Gutiérrez et al., 2003).

Currently, antivenoms are produced in three formats: IgGs (150 kDa), and their derivative F(ab’)_2_ (100 kDa) and Fab (50 kDa) fragments. Conventional snakebite antivenoms are obtained from the sera of horses and sheep which have been hyper-immunised with controlled doses of venom (León et al., 2018). Whilst generally effective at treating systemic envenomation, serum antivenoms have numerous drawbacks. These include a high risk of adverse effects, poor treatment of tissue necrosis, and low concentrations of therapeutically active antibodies (de Silva et al., 2016; Gutiérrez et al., 2003; Segura et al., 2013). There is subsequently a need to improve the design and production of antivenoms.

In recent years, recombinant protein production and *in vitro* antibody selection have diversified the range of antivenom scaffolds available for development. Nanobodies (13-15 kDa) and single chain variable fragments (scFvs - 27 kDa) are two alternative scaffolds which have been successfully selected to neutralise venom toxins (Fernandes et al., 2021; Laustsen et al., 2018). A host of alternative scaffolds could also potentially be applied to antivenom development, including affimers, affibodies, and DARPins. These have been outlined and reviewed by Jenkins et al. (2019). The alternative and conventional scaffolds together constitute a rich design field for antivenom development, spanning a wide molecular size range and variable valency configurations. Next-generation antivenoms may be further improved by affinity maturation, which can enable the production of antibodies that surpass the affinity limits of natural immune systems (Foote and Eisen, 2000). *In vitro* selection approaches can be adapted to specifically select for fast on-binding rates (by reducing the amounts of input antibody and reducing binding time) or slow off-binding rates (by increasing washing stringency and using soluble binding competitors) (Lu et al., 2003). Antibody humanisation can additionally improve the immune-tolerance of antivenom therapies, reducing the risk of side-effects (Wang et al., 2021). These technologies together have great potential to improve the efficacy and safety of antivenom therapy.

Despite the range of scaffolds that are being and could be applied to antivenom development, there is no clear consensus as to the extent that scaffold format impacts treatment efficacy. Clinical and pre-clinical comparisons of the three conventional antivenom formats have variably indicated the influence of scaffold choice on treatment outcome. Some studies report antivenom choice to have a significant impact on certain treatment outcome metrics (Boels et al., 2020; Bush et al., 2015; Ismail and Abd-Elsalam, 1998; Kurtović et al., 2021; Mascarenas et al., 2020; Morais et al., 1994; Resiere et al., 2020; Rivière et al., 1998, 1997; Wilson et al., 2022). Conversely, other studies report no impact (Carotenuto et al., 2021; Chaves et al., 2003; Dart and McNally, 2001; Gerardo et al., 2021; León et al., 2001, 1999, 1997). These studies assess different host species, venoms, antivenoms, doses, and treatment metrics, hindering a unified assessment of the impact of antivenom format. The studies additionally do not consider alternative scaffolds. Standardised comparisons of a range of different scaffolds in different treatment scenarios would be required to determine the pharmacodynamic influence of scaffold format.

Computational modelling can aid in answering this question by facilitating the assessment of antivenom pharmacokinetics and pharmacodynamics in isolation, since antivenoms with identical neutralising capacities can be simulated. These factors are difficult to control for *in vivo*, as different antivenoms may have different modes and efficacies of binding to the same toxins. Computational modelling additionally allows the standardised, systematic, and rapid assessment of a much wider area of antivenom parameter space than would be possible in vivo. Parameter sweeps can enable the identification of effective treatments which can be tested in the lab, with resulting experimental data used to re-parameterise and improve the model as required. In this way, computational modelling can elucidate the underlying system pharmacodynamics of envenomation treatment. Computation is additionally low-cost and can help reduce reliance on animal testing, by informing experimental dosing levels and enabling the pre-clinical screening of different antivenoms under different treatment scenarios.

Despite the benefits of computation, there are few existing mathematical models of venom-antivenom treatment systems. Sevcik et al. (2004) previously established a pharmacodynamic model of *Tityus discrepans* scorpion envenomation and F(ab’)_2_ antivenom treatment, however this model did not account for the elimination of antivenom or the spontaneous dissociation of venom-antivenom complexes. We subsequently defined a compartmental model of systemic snakebite envenomation and treatment to more completely describe the system and facilitate the simulation of varied venoms and antivenoms (Morris et al., 2022). This model is capable of tracking the levels of venom, a monovalent antivenom, and neutralised venom as they move through a central compartment of blood and well-perfused tissue, and a peripheral compartment of poorly-perfused tissue. We additionally defined mathematical relationships between antivenom scaffold size and three key pharmacokinetic parameters controlling elimination and tissue distribution. The model is parameterised to simulate the treatment of different venoms and antivenoms, making it an ideal testing ground to identify the parameter values and combinations that lead to improved neutralisation coverage.

In this study, we apply our monovalent binding model and define a new bivalent binding model to explore the effects of antivenom size, dose, affinity, and valency, on systemic envenomation treatment. We simulate envenomation treatment at variable timepoints, using a set of 200,000 model antivenoms that vary across five design parameters. We simulate and compare the treatment of two model venoms: the elapid *Naja sumatrana* (equatorial spitting cobra), and the viper *Cryptelytrops purpureomaculatus* (mangrove pit viper). These are well-characterised venoms with differing pharmacokinetic properties, which can be used to show the general applicability of our approach and draw broad conclusions about the effective treatment of different types of venom. *N. sumatrana* venom is rich in low molecular weight neurotoxins which rapidly absorb and distribute through the body (Chong et al., 2019; Yap et al., 2014b, 2014a). Conversely, *C. purpureomaculatus* venom is a haemorrhagic venom with a longer elimination half-life and more high molecular weight toxins (Sim et al., 2013; Zainal Abidin et al., 2016). The model outputs from these simulations were assessed, and global sensitivity analysis (GSA) and global parameter optimisation were employed to define the most influential antivenom parameters and their bounds for improved treatment. This workflow is outlined in Figure 1. Guidelines for antivenom design resulting from our model are outlined in the discussion.

**Figure 1.**
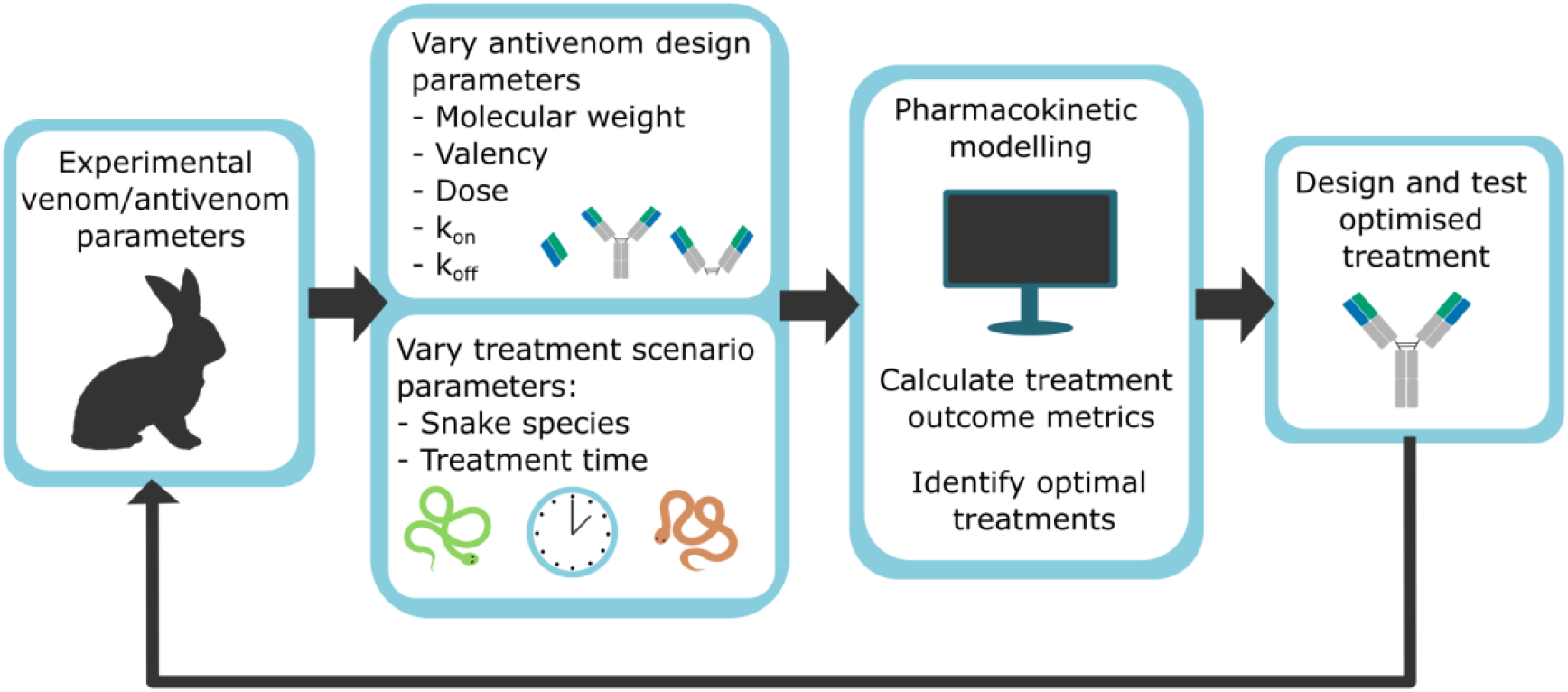
Schematic of a computational modelling workflow applied to antivenom design. The model is parameterised using experimental pharmacokinetic data. We used the model to simulate envenomation and treatment while varying several antivenom parameters and treatment scenario parameters. These simulations were used to identify antivenom parameter values and combinations for improved treatment. In future, these predicted solutions could be used to inform antivenom design and to re-parameterise the model in a positive feedback loop.

## 2 Material and Methods

### 2.1 Venom and antivenom compartmental model design

We used a two-compartment pharmacokinetic model to describe envenomation and treatment. A two-compartment structure was chosen as literature reviews have shown that this is most commonly used to describe the *in vivo* dynamics of both venoms and antivenoms, and thus offers a good representation of envenomation-treatment system dynamics (Morris et al., 2022; Sanhajariya et al., 2018). To simulate an intramuscular injection, venom is introduced to the central compartment at a set absorption rate (k_a_) and limited bioavailability (F). After a time delay, antivenom is injected to the central compartment as an instantaneous, intravenous (I.V) bolus dose with 100% bioavailability. Antivenom can bind and neutralise venom at a rate of k_on_, and spontaneous dissociation of venom-antivenom complexes is accounted for with a rate of k_off_. Venom, antivenom, and neutralised venom can move between the central and peripheral compartment at individual intercompartmental transfer rates (k_12_ and k_21_), and are removed from the central compartment at individual elimination rates (k_10_). The model parameters and their units are listed in Table 1. Antivenom k_on_ and k_off_ units were transformed to ml.ng^-1^h^-1^ and h^-1^ respectively prior to simulation, using antivenom molecular weight. All simulations in this paper use a maximum runtime of 30 hours.

**Table 1.** Pharmacokinetic parameters of the envenomation and treatment model

| Parameter | Units | Description |
| --- | --- | --- |
| $k_{10}$ | $h^{-1}$ | Elimination rate constant |
| $k_{12}$ | $h^{-1}$ | Transfer rate constant from central to peripheral compartment |
| $k_{21}$ | $h^{-1}$ | Transfer rate constant from peripheral to central compartment |
| $k_a$ | $h^{-1}$ | Intramuscular absorption rate constant |
| $k_{on}$ | $M^{-1}s^{-1}$ | Antivenom on-binding rate constant |
| $k_{off}$ | $s^{-1}$ | Antivenom off-binding rate constant |
| $V_c$ | ml | Volume of central compartment |
| $V_p$ | ml | Volume of peripheral compartment |
| $F$ | % | Intramuscular bioavailability |

### 2.2 Envenomation-Treatment Model Equations

Envenomation and treatment were modelled using ordinary differential equations (ODEs), which describe the rate of change in concentration of each species in each compartment over time. We used a monovalent binding model which we previously defined (Morris et al., 2022), alongside a new bivalent binding model. The monovalent model was applied as described in our previous paper, with the exception that in this study we estimated the pharmacokinetic rate parameters of neutralised venom-antivenom complexes using particle size (instead of assuming these to be equal to the antivenom’s parameters).

For both the monovalent and bivalent binding case, Eqs. 1a-d were used to describe the movement of venom around the body following an intramuscular bite, prior to antivenom treatment. In all model equations, the number 1 denotes the intramuscular injection compartment, 2 the central compartment, and 3 the peripheral compartment. The antivenom is referred to as A, T refers to toxins, and E to eliminated toxins. Monovalent or bivalent antivenoms with one toxin bound are referred to with N, and bivalent antivenoms with two toxins bound are referred to with NN. Subscript letters signify the species that the rate parameter is associated with. The intramuscular venom injection compartment (T1) is tracked by mass as its volume cannot be defined (Eq. 1a).

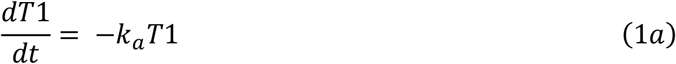

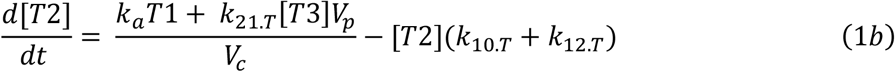

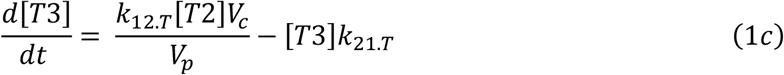

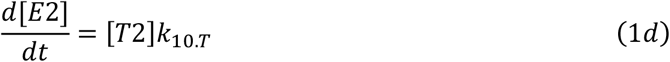

The monovalent binding model is shown in Eqs 2a-g. These equations were adapted to account for bivalent binding using bivalent binding terms (BBTs). In the monovalent binding case, all BBTs = 0.

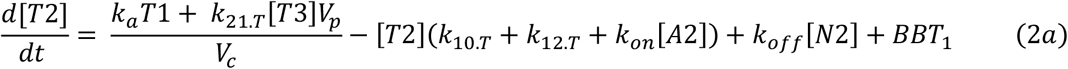

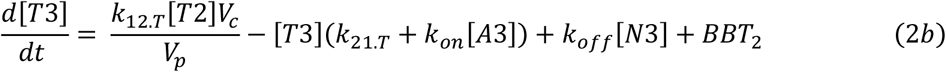

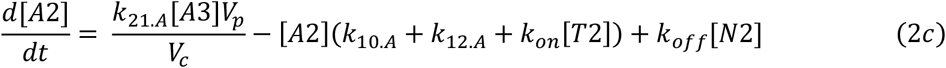

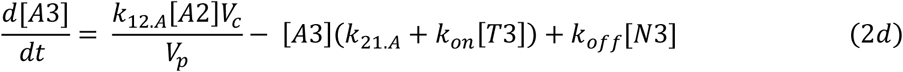

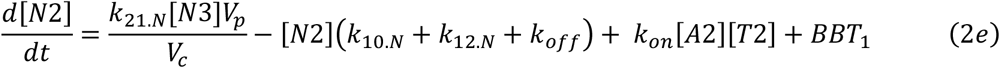

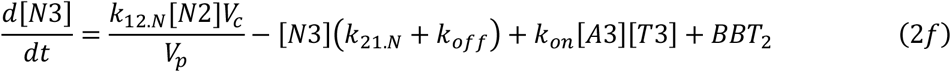

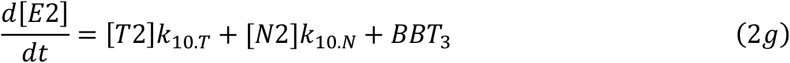

The bivalent toxin binding model allows the neutralisation of up to two toxins per antivenom particle. We assumed that the k_on_ and k_off_ rates for both binding reactions were the same. To convert the monovalent binding model to a bivalent binding model, we used the same ODE system as in Eqs. 2a-g and set the BBTs to the corresponding terms listed in Eqs. 3a-c. The bivalent antivenom system additionally included Eqs. 4a-b, which describe the dynamics of doubly bound antivenom particles in the central and peripheral compartment, respectively.

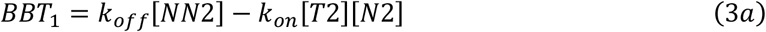

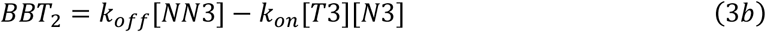

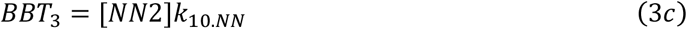

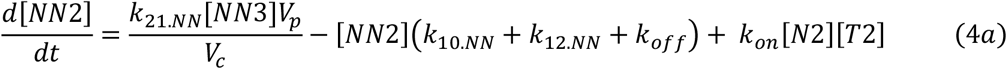

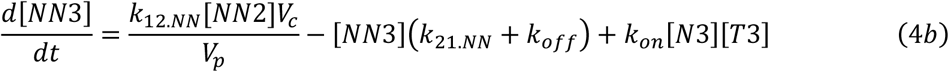

To run the simulations, envenomation was first modelled with Eqs. 1a-d, after which antivenom was introduced and the equation set was switched to Eq. 1a and either the monovalent or bivalent binding system. ODEs were solved numerically in Python: the equations prior to treatment were solved using the Runge-Kutta method of order 5(4), and the equations following treatment were solved using the Radau solver for stiff ODEs.

### 2.3 Pharmacokinetic parameters

We parameterised our model using experimental data from rabbits, as our previous literature review of experimental studies revealed rabbits to be the most commonly used pharmacokinetic model system for both venom and antivenom (Morris et al., 2022). The rabbit system has the benefits that it facilitates *in vivo* validation in the form of rescue experiments, it is large enough to enable blood sampling over days to track longer envenoming timecourses, and we can apply the parameters of real toxins and venoms. All simulations in this paper assume a rabbit weight of 2 kg. Venom pharmacokinetic parameters were taken from existing experimental studies of *N. sumatrana* and *C. purpureomaculatus* envenomation (Table 2) (Sim et al., 2013; Yap et al., 2014b). The central and peripheral compartment volumes in the model were taken to be those of the venom.

**Table 2.** Venom pharmacokinetic parameters used in this study. Compartment volumes have been standardised by body weight (BW), using the midpoint of reported rabbit weight ranges.

| Snake species | Parameter | Value | Source |
| --- | --- | --- | --- |
| *N. sumatrana* | Molecular weight (kDa) | 9 | Weighted average of proteomic venom composition (Yap et al., 2014a) |
| | $k_a$ ($h^{-1}$) | 2.2 | Approximated (Morris et al., 2022) |
| | $k_{10}$ ($h^{-1}$) | 0.0948 | Calculated using $CL/V_c$ (Yap et al., 2014b) |
| | $k_{12}$ ($h^{-1}$) | 0.4 | Yap et al., 2014b |
| | $k_{21}$ ($h^{-1}$) | 0.5 | Yap et al., 2014b |
|  | F (%) | 41.9 | Yap et al., 2014b |
| | $V_c$ ($ml\ kg^{-1}$) | 500 | Yap et al., 2014b |
| | $V_p$ ($ml\ kg^{-1}$) | 400 | Yap et al., 2014b |
| *C. purpureomaculatus* | Molecular weight (kDa) | 57 | Weighted average of proteomic venom composition (Zainal Abidin et al., 2016) |
| | $k_a$ ($h^{-1}$) | 0.7 | Approximated (Morris et al., 2022) |
| | $k_{10}$ ($h^{-1}$) | 0.141 | Sim et al., 2013 |
| | $k_{12}$ ($h^{-1}$) | 2.19 | Sim et al., 2013 |
| | $k_{21}$ ($h^{-1}$) | 0.48 | Sim et al., 2013 |
|  | F (%) | 41.6 | Sim et al., 2013 |
| | $V_c$ ($ml\ kg^{-1}$) | 390 | Sim et al., 2013 |
| | $V_p$ ($ml\ kg^{-1}$) | 1800 | Sim et al., 2013 |

We predicted the k_10_, k_12_, and k_21_ pharmacokinetic parameters of antivenoms and neutralised toxins from particle molecular weight, using regression relationships defined in our previous work (Eqs. 5-7) (Morris et al., 2022). In all three equations, MW refers to molecular weight in kDa. These equations together result in smaller scaffolds more rapidly and deeply perfusing into tissue and being eliminated more quickly, which is in line with the expected trend (Datta-Mannan, 2019; Li et al., 2017).

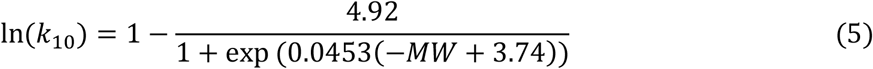

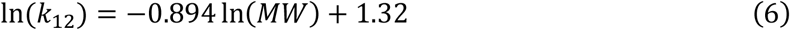

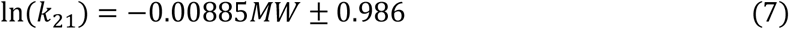

Where named scaffolds are simulated in this paper, we used our previously generated parameter sets for IgGs, F(ab’)_2_s, Fabs, scFvs, and nanobodies (Table 3) (Morris et al., 2022). The IgG k_10_ was calculated by taking a geometric mean of experimental parameters collected in a literature review, to account for the extended IgG half-life (Ober et al., 2004). Elimination half-lives (t½_β_) of scaffolds were calculated using Eqs. 8a-b, where β is the pharmacokinetic constant of the terminal phase.

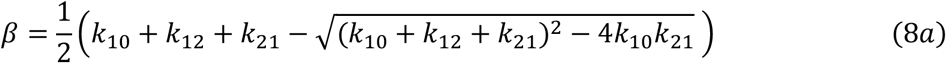

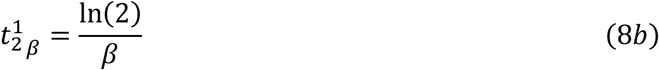

**Table 3.** Pharmacokinetic parameters of named antivenom scaffolds

| Parameter | Nanobody | scFv | Fab | F(ab')<sub>2</sub> | IgG |
| --- | --- | --- | --- | --- | --- |
| **Molecular weight (kDa)** | 15 | 27 | 50 | 100 | 150 |
| **Valency** | 1 | 1 | 1 | 2 | 2 |
| **k<sub>10</sub> (h<sup>-1</sup>)** | 0.126 | 0.0710 | 0.0341 | 0.0212 | 0.0120 |
| **k<sub>12</sub> (h<sup>-1</sup>)** | 0.334 | 0.198 | 0.114 | 0.0613 | 0.0426 |
| **k<sub>21</sub> (h<sup>-1</sup>)** | 0.327 | 0.294 | 0.240 | 0.154 | 0.0989 |

### 2.4 Venom molecular weight approximation and antivenom dose calculation

We approximated the molecular weight of the venoms to enable calculation of antivenom dose in terms of the ratio of venom particles to antivenom particles. To approximate venom molecular weight, we selected proteomics studies of *N. sumatrana* and *C. purpureomaculatus* venoms and calculated the weighted average mass of the toxin families contained in each (Yap et al., 2014a, p. 2014; Zainal Abidin et al., 2016). The details of these calculations are described in the Electronic Supplementary Materials (ESM, Section 1). *N. sumatrana* venom was simulated with an average molecular weight of 9 kDa, and *C. purpureomaculatus* venom with an average molecular weight of 57 kDa.

To calculate the mass dose of antivenom from a given ratio, we first scaled the injected venom mass by its bioavailability to obtain the mass of venom in compartmental circulation (Eq. 9a). We then converted the circulating venom mass into moles, scaled this by the required dosing ratio, and multiplied this by antivenom molecular weight to obtain the final antivenom mass (Eq. 9b). In Eqs. 9a-b, venom mass and antivenom mass are in units of mg, and venom and antivenom molecular weight (MW) are in units of Daltons. Based on these equations, in a 1:1 venom: antivenom treatment dose, the number of antivenom particles would equal the number of venom particles that could enter the central compartment.

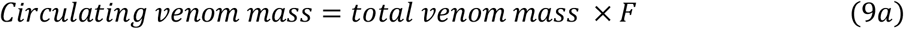

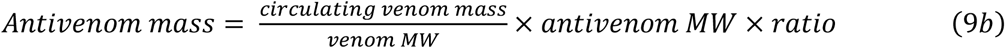

### 2.5 Definition of treatment outcome metrics

To quantify the effectiveness of different antivenom scaffolds, we defined and compared three metrics indicative of treatment outcome: the venom area under the curve (AUC), the time that the venom concentration remained over a threshold (time over threshold - TOT), and the venom AUC over a threshold (AUC-OT). The AUC metric represents the total exposure of a body compartment to venom, TOT indicates the time to reduce the venom level below a damaging concentration, and AUC-OT captures the exposure of a body compartment to a damaging venom concentration. The calculation of these metrics is visualised in the ESM (Section 2, Fig. S1). AUC metrics were calculated by the trapezoidal rule, and all metrics were calculated from the point of antivenom administration to the end of the simulation. The peripheral compartment AUC-OT metric was used in this study. Its benefits are threefold: firstly, it considers both concentration and time, unlike TOT. Secondly, it excludes negligible, persisting levels of venom that would be unlikely to cause systemic effects, unlike AUC. Finally, applying this metric to the peripheral compartment generates a wider range of output values than the central compartment, where the assumptions of homogenous mixing and instantaneous antivenom injection frequently lead to an AUC-OT of 0. This increased dynamic range makes treatment comparisons easier and more informative. These dynamics are illustrated in the ESM (Section 3).

A 50 ng/ml venom concentration was defined as an approximate threshold for systemic symptoms and was used for AUC-OT calculation in both *N. sumatrana* and *C. purpureomaculatus* envenomation. As we were unable to find data on clinical envenoming severities for *N. sumatrana* or *Cryptelytrops* family snakes, we based the threshold on experimental studies of *N. atra* and *N. kaouthia*. For comparison, the murine I.V. LD_50_ of Malaysian *N. sumatrana* snakes varies from 0.50 – 0.96 µg/g, making it less toxic than *N. atra* (0.29 µg/g, subcutaneous LD_50_) and *N. kaouthia* (0.18 µg/g, I.V LD_50_) (Liu et al., 2020; Tan et al., 2022, 2016). *C. purpureomaculatus* venoms have a similar murine I.V LD_50_ to *N. sumatrana* (0.45-0.9 µg/g) (Tan and Tan, 1988). There are several clinical accounts of *N. atra* envenomation and treatment. In one paper, blood concentrations ranging from 6.2-197.1 ng/ml coincided with systemic symptoms, with one patient reporting a 25.1 ng/ml blood concentration having presented to the clinicians shortly following envenomation (Lin et al., 2022). Another account described systemic symptoms with blood concentrations of 228 – 1270 ng/ml, and mild, non-systemic symptoms at concentrations of 0 – 21 ng/ml (Hung et al., 2003). A third account described mild symptoms at 21.2 – 93.6 ng/ml, and moderate symptoms at 147.3 ng/ml (Liu et al., 2018). Clinical gradings of *N. kaouthia* envenomation have found blood concentrations of up to 24.3 ng/ml to not result in systemic symptoms, whereas severe systemic neurotoxicity has been found at blood concentrations of 103.2 ng/ml (Faiz et al., 2017).

### 2.6 Scaffold sensitivity to dosing

We assessed the sensitivity of scaffold size to inadequate dosing. In real life, snakes inject variable and unknown amounts of venom which adds a degree of uncertainty as to effective antivenom dose levels (Tibballs, 2020; World Health Organisation, 2016). To gauge the difference in treatment outcome between minimal and maximal dosing, we calculated the fold change in AUC-OT score when venom: antivenom dosing ratios were increased from 1:1 to 1:10. To implement this, we generated pharmacokinetic parameter sets for monovalent antivenoms ranging in size from 8 to 150 kDa, at intervals of 1 kDa (Eqs. 5-7). We used each scaffold to simulate treatment of 3 mg venom at 3 hours, using venom: antivenom dosing ratios of 1:1 and 1:10. Antivenom k_on_ rates were set to 1×10^5^ M^-1^ s^-1^ and k_off_ to 1×10^−4^ s^-1^, giving a K_D_ of 1 nM which represents a high affinity antivenom. The fold-change was calculated by dividing the AUC-OT score from the 1:1 dose simulation by the 1:10 simulation.

### 2.7 Theoretical antivenom parameter sampling method

To explore the impact of different antivenom parameters on treatment outcome, we simulated treatment with a set of theoretical antivenoms and assessed the resulting AUC-OT scores. Five parameters were varied simultaneously across the theoretical antivenom parameter set: molecular size (which controlled antivenom k_10_, k_12_, and k_21_), the ratio of venom to antivenom particles, k_on_, k_off_, and valency. The upper and lower bounds of these parameters are listed in Table 4. A reasoning behind the bounds is provided in the ESM, Section 6.

**Table 4.** Minimum and maximum theoretical antivenom parameter bounds

| Parameter | Minimum | Maximum |
| --- | --- | --- |
| Antivenom dose ratio (venom: antivenom) | 1:1 | 1:10 |
| Antivenom molecular size (kDa) | 15 | 150 |
| $k_{on}$ ($M^{-1}s^{-1}$) | $1 \times 10^3$ | $1 \times 10^6$ |
| $k_{off}$ ($s^{-1}$) | $1 \times 10^{-6}$ | $1 \times 10^{-3}$ |
| Valency (categorical input) | 1 | 2 |

To generate the antivenom parameter set, we used Latin Hypercube sampling to generate 200,000 pairs of molecular weight and dose parameters. Latin Hypercube sampling is a computationally efficient way to generate randomised sample sets, with less datapoint clustering than random sampling (McKay et al., 1979). We next generated 200,000 k_on_ and k_off_ sample pairs that were evenly spaced in log space, as these parameters span several orders of magnitude. We randomly combined the k_on_ and k_off_ parameter pairs with the molecular weight and dose parameters and assigned half to monovalent and half to bivalent binding. The resulting k_on_/k_off_ combinations weight the final sample towards a K_D_ of 1×10^−8^ – 1×10^−10^ M^-1^s^-1^.

### 2.8 Theoretical antivenom treatment simulations

The theoretical antivenom parameter set was used to simulate treatment of *N. sumatrana* and *C. purpureomaculatus* envenomation. Each antivenom was simulated at treatment times ranging hourly from 1 to 10 hours, following a 3 mg intramuscular venom injection. The peripheral AUC-OT score of each simulation was recorded. To examine the impact of uncertainty in the antivenom and neutralised venom k_10_/k_12_/k_21_ parameter predictions, we additionally randomly applied variability within 2-fold to these predictions and repeated the simulations.

### 2.9 Global parameter optimisation

We analysed the theoretical antivenom simulation outputs to find the parameter combinations that led to the lowest AUC-OT scores. To do this, we used histograms to categorise the outputs according to their distributional ranking. For each treatment timepoint, we identified the antivenoms that resulted in the lowest 1% of AUC-OT scores. Antivenoms that consistently scored within the lowest 1% at every treatment timepoint were designated as the universal scaffolds. We also looked at the areas of parameter space that led to poor treatment by identifying the scaffolds that resulted in the highest 50% AUC-OT scores at every timepoint.

### 2.10 PAWN global sensitivity analysis

We utilised the PAWN GSA method to quantify the influence of the variable antivenom parameters on AUC-OT (Pianosi and Wagener, 2018, 2015). This method compares the cumulative distribution functions (CDFs) of a model’s output when given parameters are either fixed or varied. The difference between CDFs is calculated using the Kolmogorov-Smirnov (KS) test, with the final PAWN sensitivity index represented by a statistic (ie mean, median) of calculated KS values. Larger KS values indicate a greater influence of the parameter in question. The theoretical antivenom parameter set meets the prerequisite of non-correlation for accurate sensitivity index calculation: k_on_ and k_off_ are known to evolve independently of one another, and so different combinations can be equally likely (Poulsen et al., 2011).

To implement PAWN, we calculated the bootstrap median KS and its 95% confidence intervals using 100 resamples of 7500 simulations. Resampling was performed without replacement, which is recommended to reduce biases in the estimation of PAWN confidence intervals (Zadeh et al., 2017). PAWN sensitivity indices were calculated using the Python package SALib, and the PAWN tuning parameter was set to 10. A convergence test was conducted to ensure a sufficient bootstrap sample size, and to assess the impact of variation in the tuning parameter (ESM Section 9). We additionally calculated the PAWN index of a dummy parameter to determine the threshold above which a parameter could be deemed influential. The dummy was represented by generating a random number between 0-1 for each simulation (Pianosi and Wagener, 2018; Zadeh et al., 2017).

We conducted the bootstrap PAWN sensitivity test for the 10 hourly treatment time simulation sets. We also repeated the analysis on subsets of the output distributions. We calculated the sensitivity indices for antivenoms that produced AUC-OT scores below and above 300 ng.h.ml^-1^, to assess whether parameter influence changed when looking at effective or ineffective antivenoms. An AUC-OT score of 300 ng.h.ml^-1^ was chosen as the static threshold at the head of the output distributions to broadly distinguish antivenoms with some therapeutic effect from those with sub-therapeutic effect. To ensure the reproducibility of the GSA, we performed a Random Balanced Design – Fourier Amplitude Sensitivity Test (RBD-FAST) GSA which produced essentially the same parameter rankings as PAWN (ESM Section 11). We also conducted a PAWN GSA on the 2-fold varied dataset, which showed no real difference in the calculated sensitivity indices (ESM Section 10).

### 2.11 k_on_ and k_off_ sensitivity plots

We produced several plots showing the impact of k_on_ and k_off_ variation on treatment. First, we visualised how the k_on_ mode of the most effective antivenoms varied with treatment time. To do this, we produced kernel density estimation (KDE) curves of the k_on_ distribution of effective antivenoms at each treatment time, where the effective antivenoms were identified as those producing the lowest 1 % AUC-OT scores. The peaks of the KDE curves were extracted to obtain an approximate k_on_ mode, and the modes were plotted against treatment time. We also conducted a local sensitivity analysis (LSA) of k_on_ and k_off_, to assess the impact of independently varying these parameters. In log space, we generated 10,000 evenly spaced k_on_ values between 1×10^3^ – 1×10^6^ M^-1^s^-1^, and 10,000 evenly spaced k_off_ values between 1×10^−6^ – 1×10^−3^ s^1^. When variable k_on_ parameters were used, k_off_ was fixed to 1×10^−5^ s^-1^. When variable k_off_ parameters were used, k_on_ was fixed to 1×10^5^ M^-1^s^-1^. We used these affinities to simulate treatment of a 3 mg intramuscular dose of either *N. sumatrana* or *C. purpureomaculatus* venom. The antivenom was set to a monovalent 50 kDa scaffold, and the venom: antivenom dosing ratio was 1:5. The resulting AUC-OT scores were plotted against the varied affinity parameter.

### 2.12 Software and computing facilities

Figure 1 was produced in Inkscape. All other figures were produced in Python. All simulations were coded in Python, using JupyterLab 3.3.2. Specific packages used were NumPy 1.21.5, matplotlib 3.5.1, pandas 1.4.2, SALib 1.4.5, scipy 1.7.3, seaborn 0.11.2, and tqdm 4.64.0. The global antivenom optimisation simulations were performed using the computational facilities of the Advanced Computing Research Centre, University of Bristol - http://www.bristol.ac.uk/acrc/. Python scripts for this project are available at https://bitbucket.org/hauertlab/antivenom_optimisation/src/main/.

## 3 Results

### 3.1 Viper and elapid envenomation-treatment simulations

We expanded our previous model of systemic envenomation and antivenom treatment to enable bivalent toxin binding, and applied these models to explore the antivenom design space. The pharmacokinetic model simulates an intramuscular bite treated with intravenous antivenom, and follows the movement of venom, antivenom and neutralised venom around a central and peripheral compartment. It allows user-control of various parameters, including venom type and dose, antivenom type and dose, treatment time, and antivenom affinity. The model can be used to compare the effect of different types of antivenom therapy in different scenarios. Through this study, *N. sumatrana* envenomation was designated as a model elapid snakebite, and *C. purpureomaculatus* envenomation as a model viper snakebite. Example simulations of the elapid venom treated with a monovalent antivenom, and the viper venom treated with a bivalent antivenom, are shown in Figure 2. The antivenoms are administered at different times and dose levels, and with different k_on_ and k_off_ values to illustrate the flexibility of the model. Both antivenoms in these simulations have a high affinity with a K_D_ of 1 nM, and are capable of inducing the rapid neutralisation of circulating venom in both compartments.

**Figure 2.**
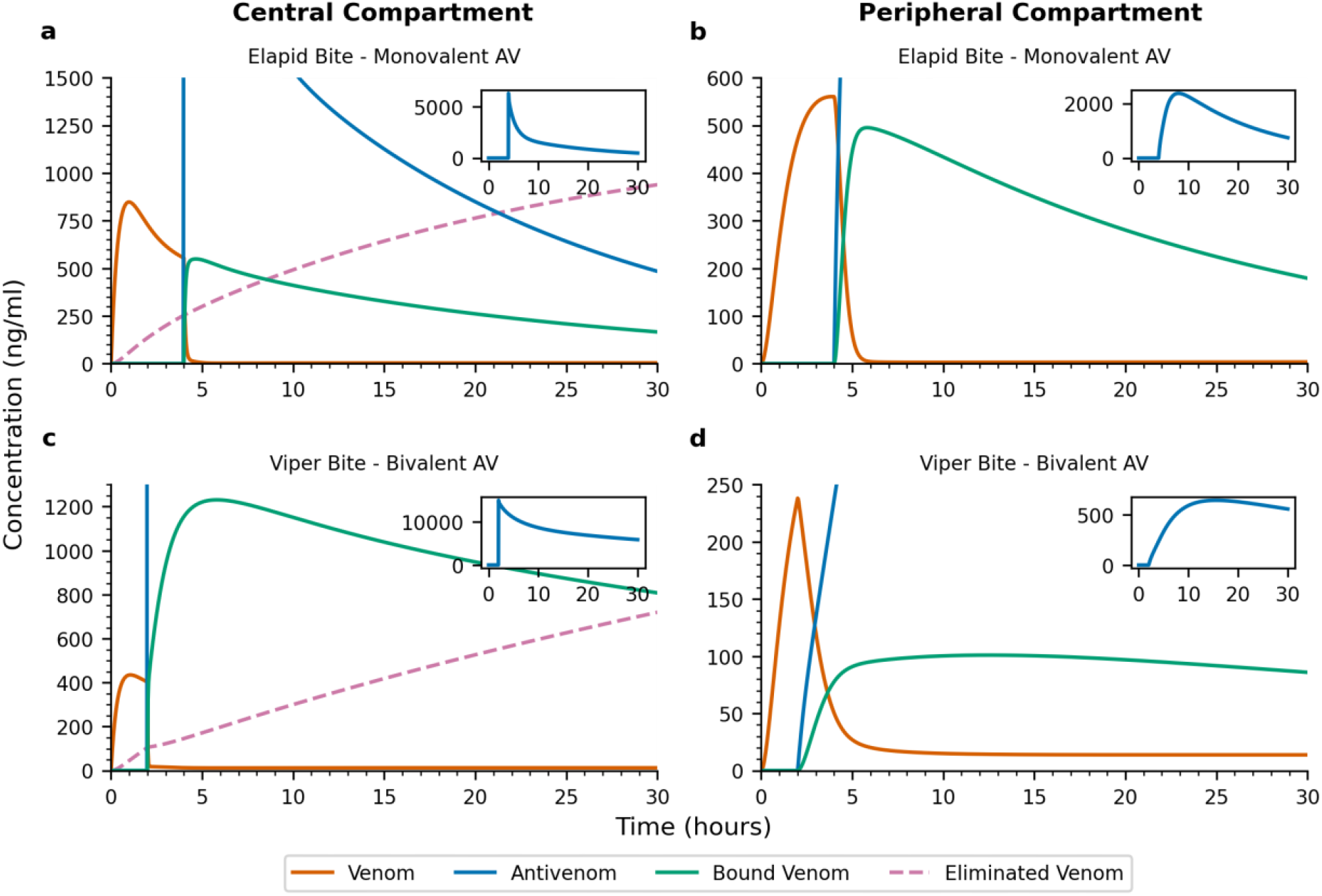
Example elapid and viper envenomation treatment simulations with high affinity monovalent and bivalent antivenoms. Both antivenoms induce rapid, extensive, and sustained toxin neutralisation. Panels on the left show the central compartment, and panels on the right show the peripheral compartment. Panels a-b show the treatment of 3 mg elapid venom at 4 hours with 6.29 mg of a monovalent nanobody antivenom, equating to 1:3 venom: antivenom ratio. The antivenom has k_on_ = 1×10^4^ M^-1^ s^-1^ and k_off_ = 1×10^−5^ s^-1^. Panels c-d show the treatment of 4 mg viper venom at 2 hours with 11.7 mg of bivalent F(ab’)_2_ antivenom, equating to a 1:4 venom: antivenom particle ratio. The antivenom has k_on_ = 1×10^5^ M^-1^ s^-1^ and k_off_ = 1×10^−4^ s^-1^. AV = antivenom.

### 3.2 At sufficient doses, low molecular weight antivenoms enhance toxin neutralisation in the tissue

We defined the peripheral compartment AUC-OT metric to represent the body’s exposure to damaging venom concentrations, enabling us to quantitatively compare different treatment modalities. We first used this to compare the performance of different antivenoms at different dosing ratios. Figure 3 shows the AUC-OT scores following treatment of elapid and viper envenomation with five different antivenom scaffolds, at dosing ratios of 1:1 and 1:10. Full profiles of the AUC-OT score evolution as the dosing ratios are increased stepwise from 1:1 to 1:10 are shown in the ESM (Section 4). Across the simulations, higher doses are required for low molecular weight antivenoms to outperform high molecular weight scaffolds due to their shorter elimination half-life. This effect is more pronounced for the viper venom, due to the venom’s longer absorption time and half-life. When dosed sufficiently, smaller scaffolds lead to lower AUC-OT scores compared to larger scaffolds at the same dosing ratio, indicating enhanced toxin clearance within the tissue. We additionally performed an analysis of central compartment AUC scores, finding that larger antivenom scaffolds outperform smaller scaffolds in the clearance of systemic blood toxins, both at minimal and saturating dose levels (ESM, Section 5).

**Figure 3.**
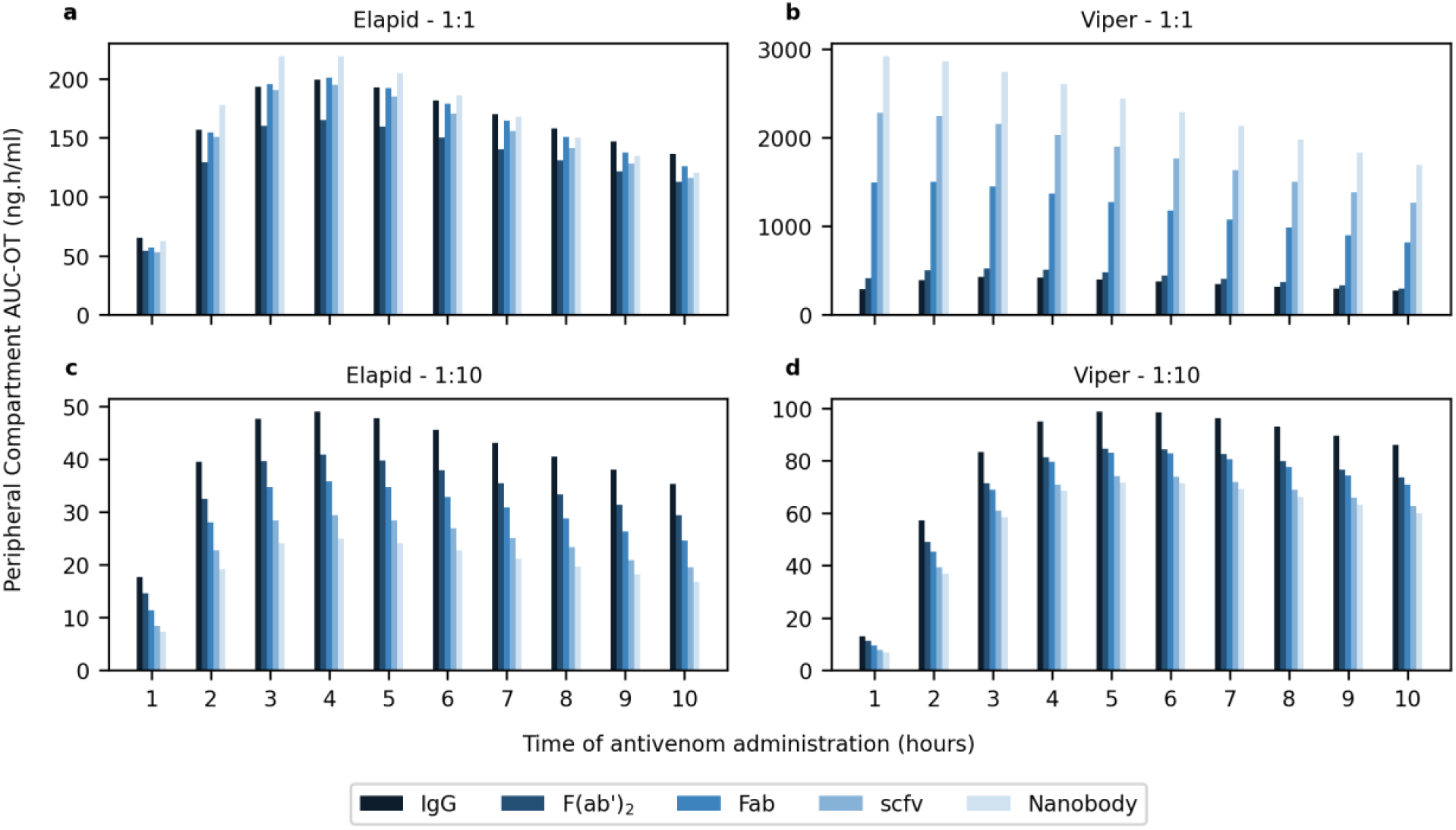
Peripheral compartment AUC-OT scores at variable dosing ratios and treatment times following elapid or viper envenomation. These plots show that provided a sufficient dose, low molecular weight antivenoms improve toxin neutralisation in the tissue. Panels a and c show the AUC-OT scores following elapid envenomation at venom: antivenom dosing ratios of 1:1 and 1:10, respectively. Panels b and d show the AUC-OT scores following viper envenomation at dosing ratios of 1:1 and 1:10, respectively. In all simulations, k_on_ = 6×10^5^ M^-1^ s^-1^ and k_off_ = 1×10^−3^ s^-1^.

### 3.3 Antivenoms which are smaller than their toxin targets may require closer clinical monitoring

We next tested the sensitivity of antivenoms of different sizes to insufficient dosing, which may be an issue in field bites given that snakes inject unknown amounts of venom (World Health Organisation, 2016). To quantify this, we calculated the fold-change in AUC-OT for antivenoms of different sizes, when dosing was varied from a ratio of 1:1 to 1:10. Larger fold-changes indicate a bigger AUC-OT shift when treating with minimal versus saturating antivenom doses, and potentially poorer treatment outcomes if the initial antivenom dose is insufficient. The fold-change plots reveal changing patterns of dosing sensitivity depending on the venom being treated (Fig. 4). The viper venom is prone to large fold-changes with smaller antivenoms, with the curve stabilising once antivenom scaffolds reach molecular sizes closer to the average molecular weight of the venom toxins (57 kDa). Conversely, the elapid curve stabilises at a much lower molecular weight owing to the smaller 9 kDa average size of the venom toxins. The average size of the toxin relative to the size of the antivenom thus seems to mediate this sensitivity, indicating that systemic viper envenomation may be more reliably treated using larger antivenom scaffolds. In a clinical context, this may mean that treatment of viper envenomation with a low molecular weight antivenom may require closer monitoring of patient symptoms and recovery. Patients may additionally be more likely to require repeated antivenom dosing. An increased requirement for redosing with Fab antivenoms compared to F(ab’)_2_ antivenoms has been clinically reported for rattlesnake envenomation (Mascarenas et al., 2020).

**Figure 4.**
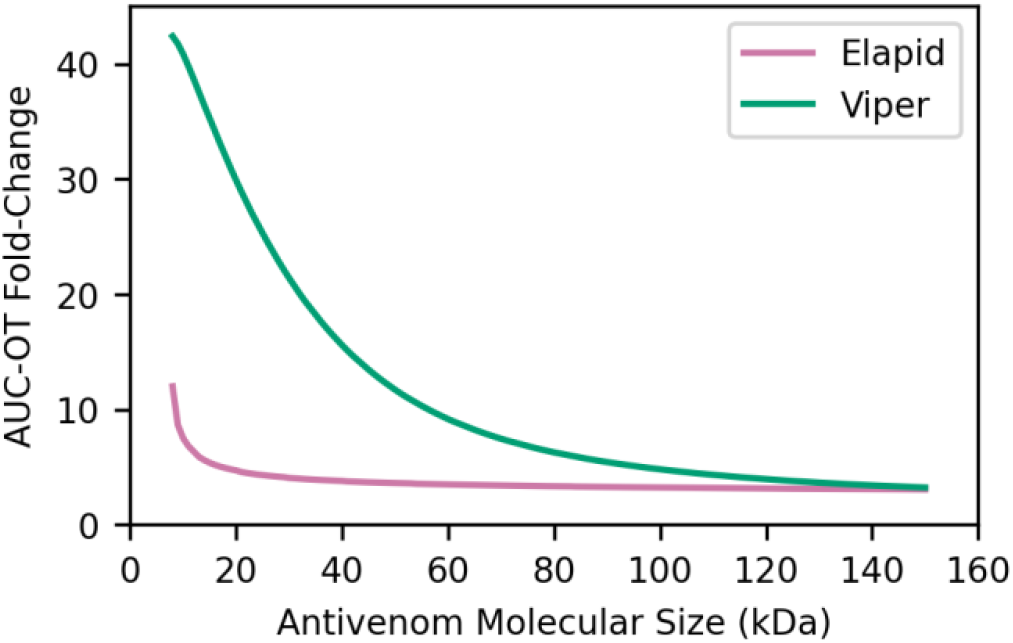
Effect of antivenom scaffold size on AUC-OT fold-change when antivenom dose is increased from 1:1 to 1:10. The viper venom is treated more reliably with high molecular weight antivenoms, whereas the elapid venom tolerates a wider range of scaffold sizes. Simulations assumed a 3 mg intramuscular venom bite, with antivenom administered 3 hours post-envenomation. k_on_ = 1×10^5^ M^-1^ s^-1^ and k_off_ = 1×10^−4^ s^-1^.

### 3.4 Effective antivenoms can span a wide area of parameter space

We next assessed the impact of simultaneously varying five antivenom parameters on treatment outcome, to define the characteristics of pharmacodynamically effective antivenoms. We used a set of 200,000 theoretical antivenoms which varied across molecular weight, antivenom dosing ratio, k_on_, k_off_, and valency. This was used to simulate treatment of elapid and viper envenomation, with simulations repeated for treatment times ranging from 1 to 10 hours post-bite. For each simulation, we recorded the AUC-OT. Figure 5 shows the AUC-OT output distribution from the elapid 3 hour treatment case. This distribution, like all of the simulated output distributions, is heavily right-skewed. This indicates that a majority of the theoretical antivenoms lead to low AUC-OT scores, and thus a wide area of the tested antivenom parameter space is conducive to effective treatment. The output distributions of all simulations are shown in the ESM (Section 7). Increasing treatment time delays result in less-skewed output distributions, reflecting a narrowing of the parameter bounds for effective treatment and a reduced maximal AUC-OT.

**Figure 5.**
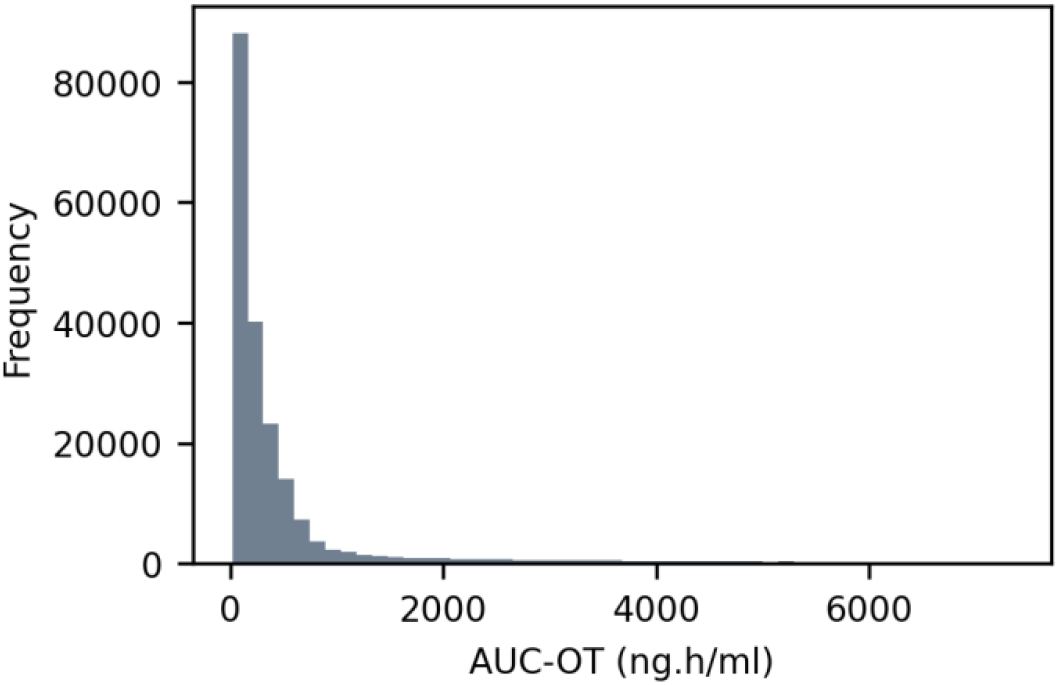
Histogram of AUC-OT scores from treatment of elapid envenomation with 200,000 theoretical antivenoms. Antivenom administered 3 hours post-bite. The right-skewed distribution illustrates that the majority of the tested antivenoms result in effective toxin neutralisation. Bin number = 50.

### 3.5 Viper envenomation confers tighter constraints on effective antivenom design

We looked at the parameter combinations of the best-performing antivenoms. Figure 6 shows three-dimensional plots of the antivenoms which result in the 1% lowest AUC-OT scores in every treatment time simulation, which we will refer to through the rest of this paper as universal scaffolds. There are 37,967 universal scaffolds for the elapid, but only 5,541 for the viper, indicating the narrower region of parameter space for effective viper envenomation treatment. The antivenoms were split into monovalent and bivalent scaffolds: 53.8% of universal elapid treatments were bivalent, compared to 76.6% of viper treatments. Across these scaffolds, monovalent binding coincides with increased baseline dose and affinity levels. Across both venoms, several general trends are apparent: universal scaffolds can be found across the entire molecular weight range of 15 – 150 kDa, however smaller antivenoms permit lower affinities and lower doses. Higher molecular weight scaffolds have visibly higher minimal effective dosing ratios. This is likely because higher doses increase the speed with which antivenom can move from the central to the peripheral compartment, counteracting the slower distribution rate for larger scaffolds. The universal scaffolds have different minimal effective doses for each snake (roughly 1:2 venom: antivenom for the elapid, and 1:5 venom: antivenom for the viper). Elapids can also tolerate a weaker affinity antibody, with the weakest K_D_s of the universal scaffolds in the order of around 1×10^−8^ M^-1^s^-1^. Conversely, the weakest K_D_ of the viper scaffolds is closer to 1×10^−9^ M^-1^s^-1^.

**Figure 6.**
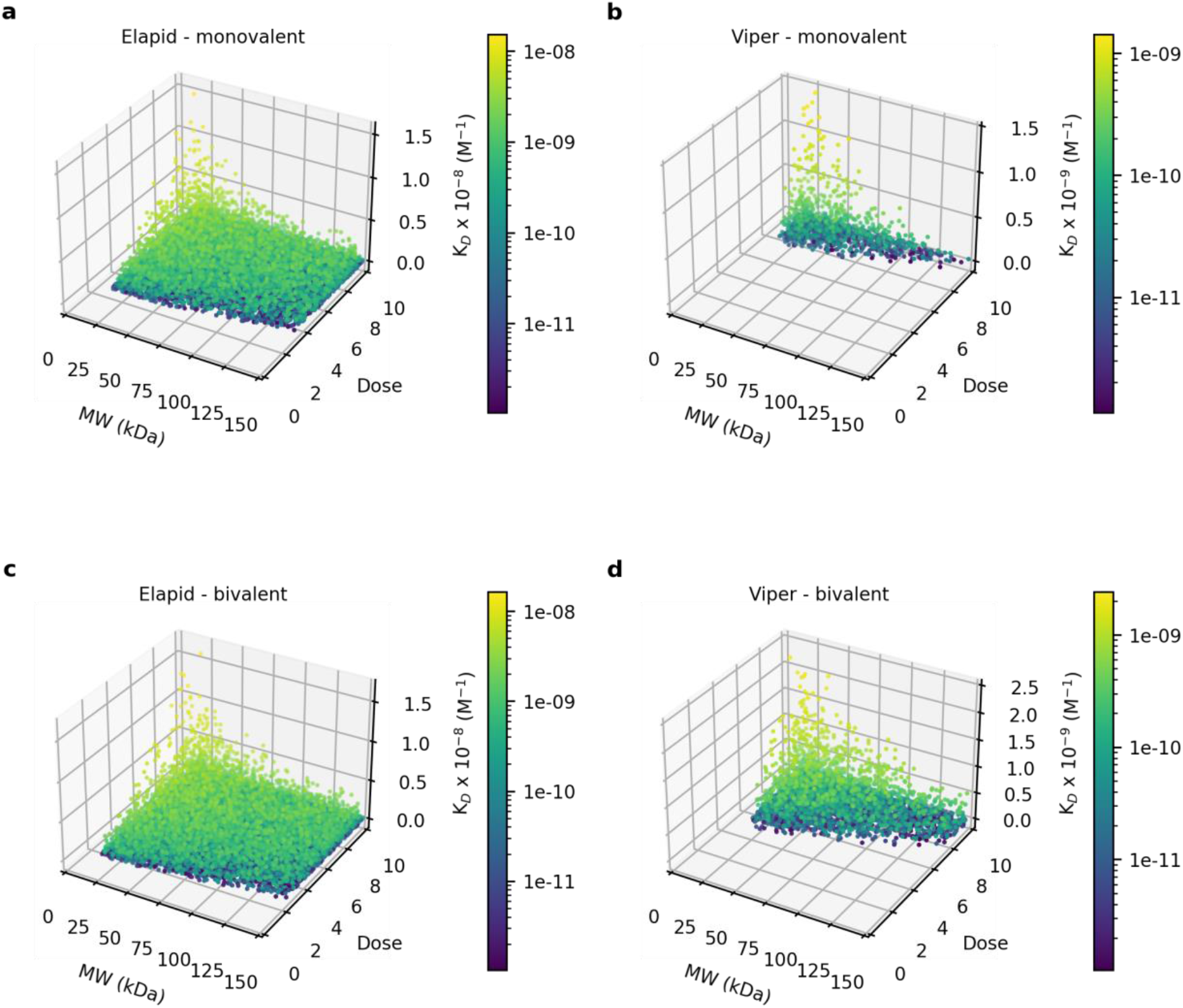
Parameter combinations of the universal antivenom scaffolds, categorised by valency. These scaffolds performed in the top 1% of theoretical antivenoms across all treatment delay simulations. Panels on the left show the elapid envenomation case, and panels on the right show the viper envenomation case. Panels a and b show the monovalent scaffolds, and panels c and d show the bivalent scaffolds. The colour bar corresponds to antivenom K_D_ (K_D_ = k_off_/k_on_). Both venom types show a tolerance for scaffolds of different sizes, and a preference for high dose and high affinity. Effective viper antivenoms however have more constrained parameter ranges.

### 3.6 Low molecular weight scaffolds with high dose and affinity are preferred for improved performance

Figure 7 shows KDE plots of the antivenom parameter combinations of the universal scaffolds. Deeper-hued contours indicate a higher probability that an antivenom was found within that area of parameter space. These plots indicate a preference for low molecular weight scaffolds, with the highest density of datapoints found below 50 kDa for both snakes. Low molecular weight scaffolds can tolerate lower dosing ratios for the same grade of treatment. The KDE plots of k_on_ against k_off_ show that highly effective elapid solutions occupy a much larger area of parameter space, indicating less stringent affinity requirements for elapid envenomation treatment. For both venoms, the highest density occurs across quite a narrow k_on_ range (10^5^ – 10^6^ M^-1^s^-1^), with a wider k_off_ range (10^−6^ – 10^−4^ s^-1^) tolerated.

**Figure 7.**
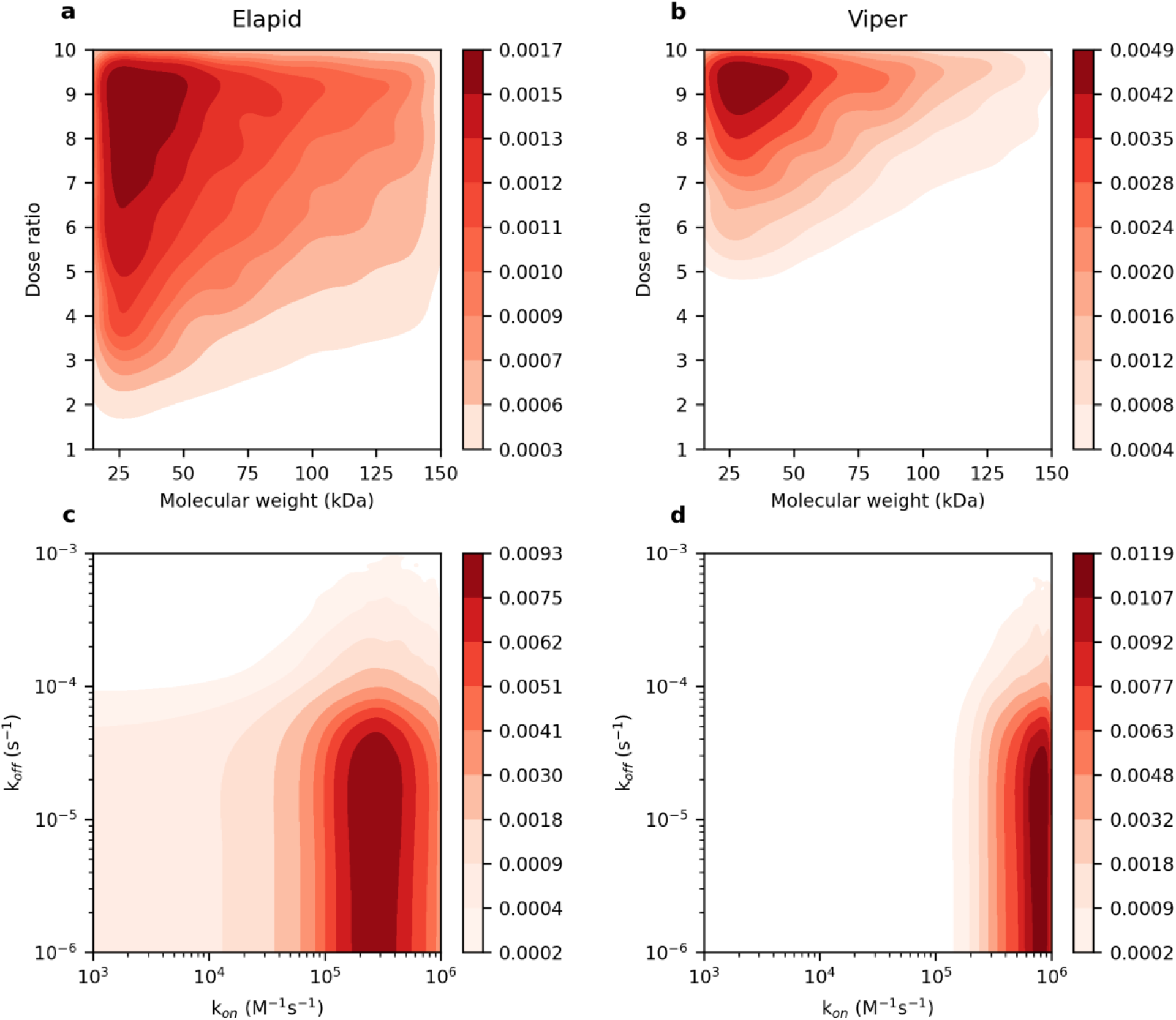
KDE plot of the universal scaffold parameter combinations. Panels on the left show top-performing elapid treatments. Panels on the right show top-performing viper treatments. The most effective antivenoms tend to have a low molecular weight, high dose, and high k_on_. Colour bars show the KDE.

We also visualised the parameters of top-performing scaffolds at specific early and late treatment times. This showed that delayed treatment leads to an increasing preference for low molecular weight scaffolds, with high dose and high affinity (ESM, Section 8). To assess the impact of uncertainty in our pharmacokinetic parameter prediction, we ran the theoretical antivenom simulations with random 2-fold variation applied to the calculated antivenom and neutralised venom parameters. The parameter space of these simulations was plotted and found to be largely unchanged (ESM, Section 10).

### 3.7 Poorly performing antivenoms are chiefly characterised by low k_on_ rates

We next visualised the parameter space of poorly performing antivenoms. Figure 8 shows a KDE plot of the parameters of antivenoms which resulted in the highest 50% of AUC-OT scores, at every treatment timepoint. Scaffolds in the bottom 50% can be found across the complete molecular size range. There is a tendency for poorer-performing scaffolds to have a low venom: antivenom dose ratio, with the highest density between 1:1 - 1:2. Affinity has the clearest impact on poor treatment, with this parameter’s density being visually distinct from that of the universal scaffolds (Figure 7). k_on_ appears to most strongly associate with poor performance, with these scaffolds nearly all having a k_on_ of below 1×10^4^ M^-1^s^-1^. Poorly performing antivenoms were found across the entire sampled k_off_ range, however for the elapid this density was mostly focused between 10^−4^ – 10^−3^ s^-1^, and for the viper this density was focused between 10^−5^ – 10^−4^ s^-1^.

**Figure 8.**
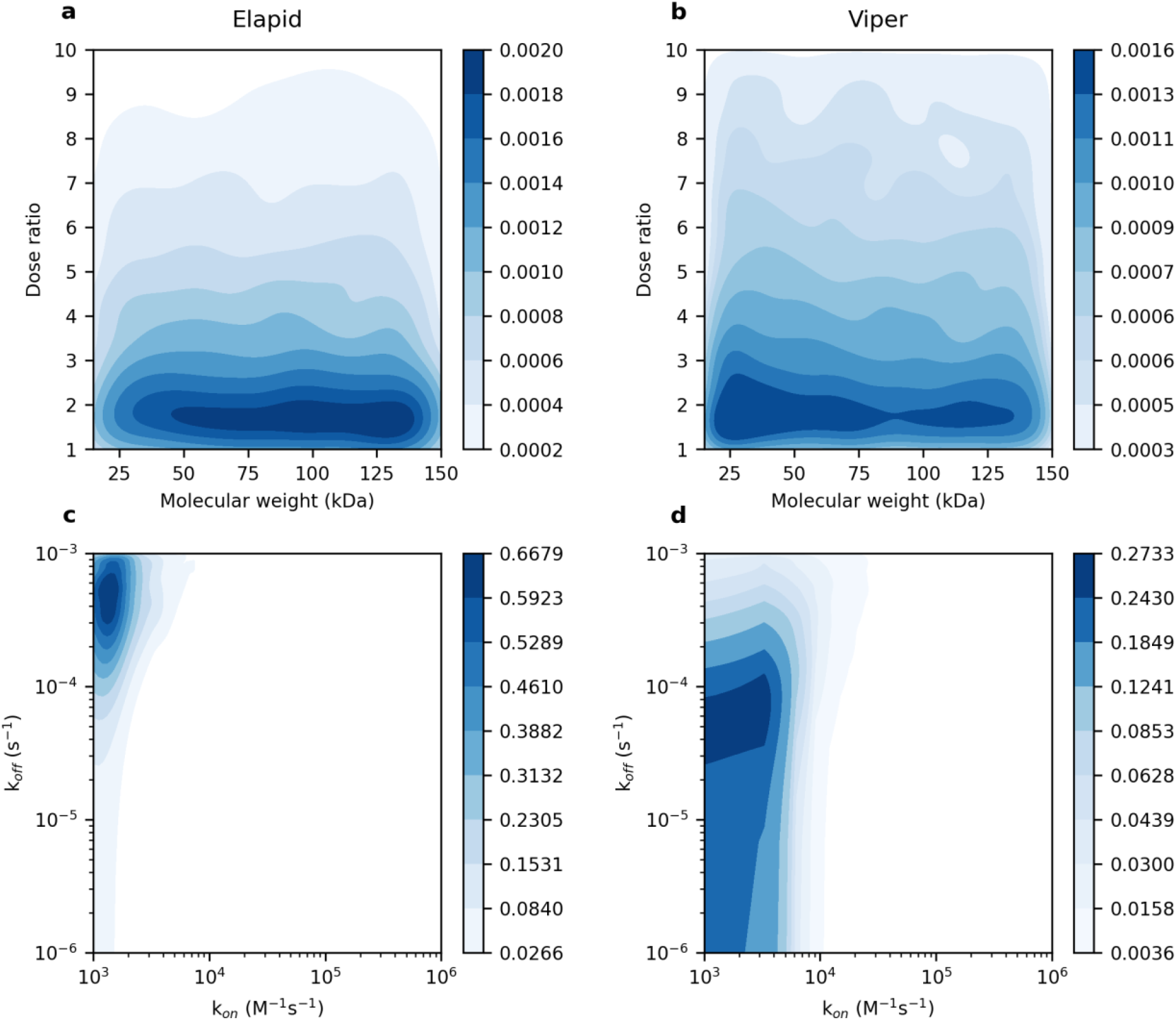
KDE plot of antivenom parameter combinations which result in the highest 50% AUC-OT scores, at every treatment time. Panels on the left show worst-performing elapid treatments. Panels on the right show worst-performing viper treatments. The density plots show the least effective treatments to generally have a low dose and low k_on_. Colour bars show the KDE.

### 3.8 Low molecular weight scaffolds have the most flexible design parameter ranges for improved performance

To identify existing antibody formats within the universal scaffold design space, we generated violin plots showing the distribution of the universal scaffold molecular weights, categorised by valency (Fig. 9). Both snake species show a preference for smaller scaffolds, with the highest density of solutions occurring between 20 – 30 kDa, of a similar size to a monovalent scFv or a bivalent nanobody (generated by connecting two nanobodies with a linker). Based on our pharmacokinetic parameter predictions, particles in this size range have an elimination half-life of 14 – 19 hours in rabbits. Scaffolds with the lowest molecular sizes have a reduced density owing to their increased elimination rates, with a noticeable decline below the approximate 25 kDa density peak. As molecular size increases, the probability density of the distribution decreases, indicating that larger scaffolds are more difficult to design for effective treatment. Viper envenomation shows this effect much more dramatically, with very limited density at the upper molecular weight ranges. This constriction is caused by treatment delays, which impart a preference for rapidly-perfusing low molecular weight binders (ESM, Section 8). Small scaffolds subsequently offer the most flexible design space for effective performance in both early and late treatment scenarios.

**Figure 9.**
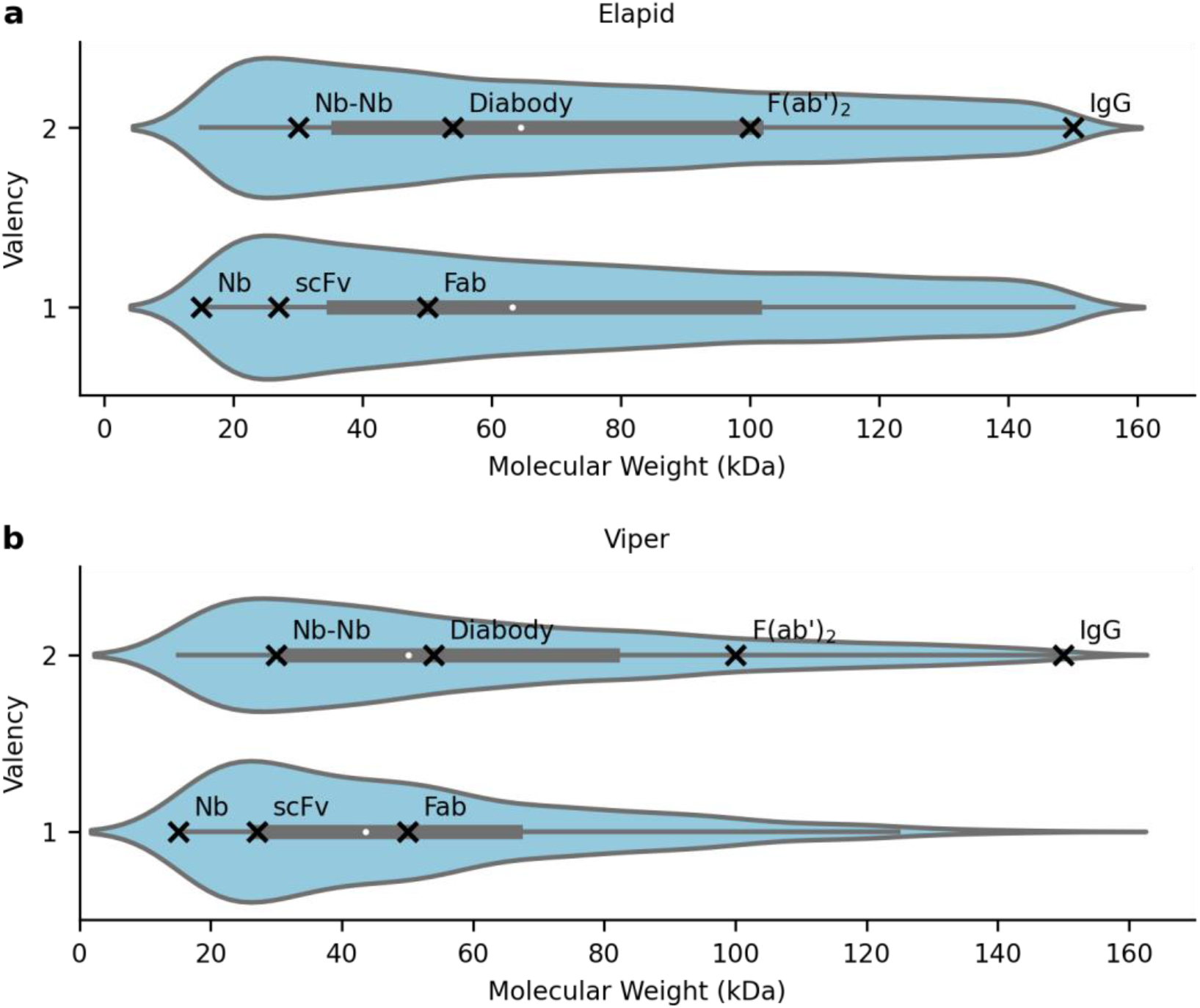
Violin plot of universal scaffold molecular weights, categorised by valency. The blue curves represent a probability distribution, with wider regions indicating a greater number of antivenoms. The greatest density occurred at low molecular weights, suggesting that these scaffolds offered enhanced flexibility around the other design parameters. Named scaffolds were plotted as follows: nanobody (Nb – 15 kDa), scFv (27 kDa), Fab (50 kDa), F(ab’)_2_ (100 kDa), IgG (150 kDa), nanobody dimer (Nb-Nb – 30 kDa), Diabody (scFv dimer - 54 kDa (Lüdel et al., 2019)).

### 3.9 Global sensitivity analysis demonstrates a conditional influence of k_on_ and k_off_ on antivenom efficacy

Global sensitivity analysis is a way to quantify the influence of different input parameters on a model’s output. It can be used to rank parameters in order of their influence and determine non-influential parameters. We conducted a PAWN GSA for each snake at each antivenom administration timepoint, to explore how parameter influence changes with early and delayed treatment times. Figs. 10a-b show the results of the GSA applied to the complete AUC-OT output distribution at each timepoint. For both the elapid and the viper, k_on_ has the largest PAWN index, and this pattern of sensitivity is relatively stable for the parameters across all treatment times. Dose, k_off_, and molecular weight have similar and low rankings, with molecular weight being the least influential parameter. Valency is below the dummy parameter threshold, indicating that it does not significantly impact treatment outcome.

**Figure 10.**
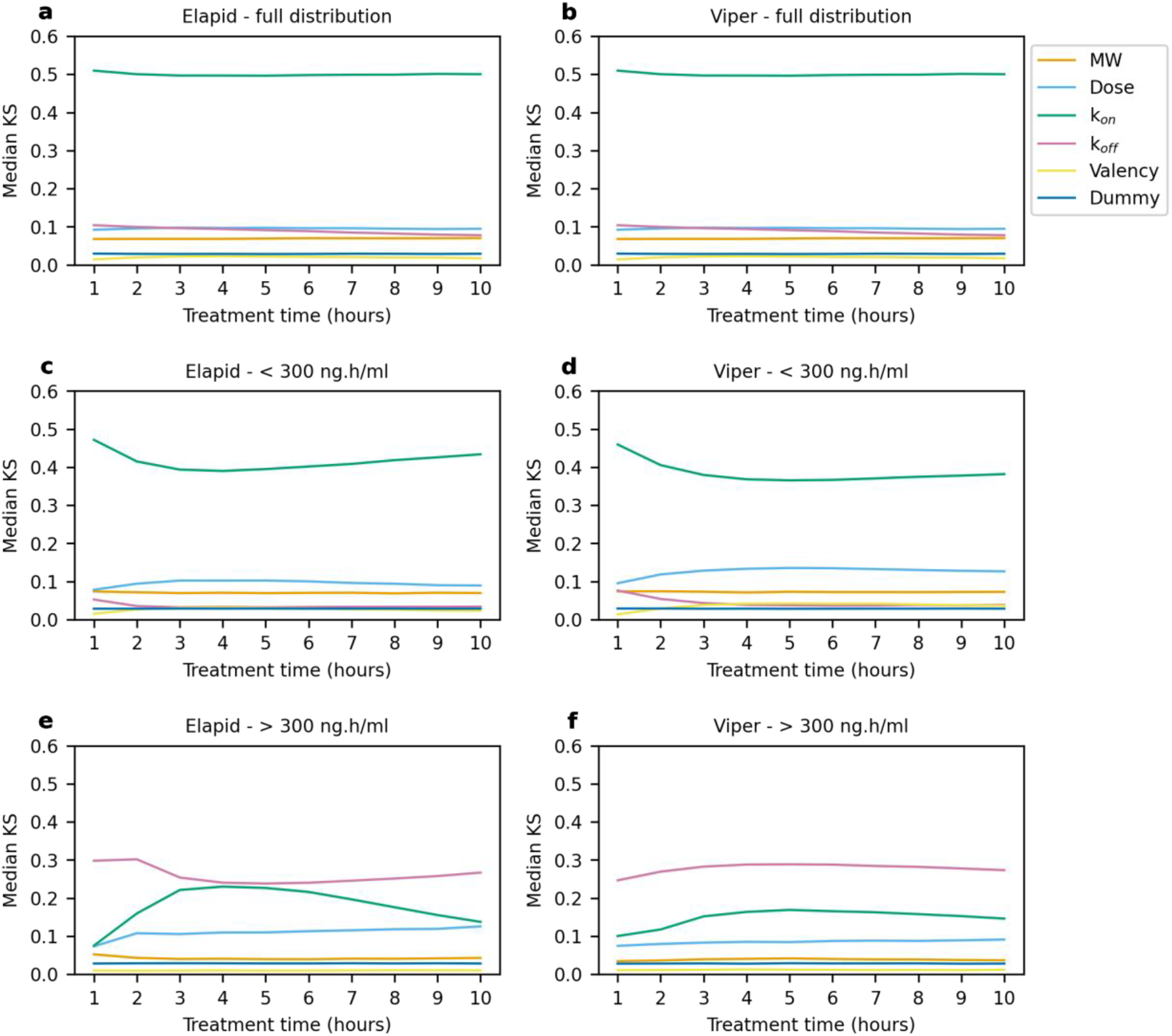
PAWN GSA showing the effect of five antivenom design parameters on AUC-OT at variable antivenom administration times. Panels a and b show PAWN applied to the complete output distribution. Panels c and d show PAWN applied to model outputs below 300 ng.h.ml^-1^, at the head of the distribution. Panels e and f show PAWN applied to model outputs above 300 ng.h.ml^-1^, at the tail of the distribution. While k_on_ is the most influential parameter overall, k_off_ is the most influential parameter for weakly effective scaffolds. The dummy parameter sets a threshold of non-influence. 95% confidence intervals were plotted as clouds around line graphs.

We next performed the PAWN GSA on AUC-OT values below or above 300 ng.h/ml, to assess parameter influence on therapeutically active or sub-therapeutic antivenoms, respectively. Slicing the distribution in this way results in noticeable changes to the resulting PAWN sensitivity indices. Panels c and d show the PAWN indices for scaffolds which are already having some therapeutic effect. These indices are largely similar to those of the complete distribution, which is to be expected as all of the output distributions are right-skewed. k_on_ remains the most influential parameter, however there is noticeably more time-dependent variation. In this GSA, dose becomes the second most influential parameter, followed by molecular weight. k_off_ overlaps closely with the dummy threshold indicating a negligible effect on the AUC-OT metric. Panels e and f show the PAWN indices for scaffolds with a more limited or absent therapeutic effect. In these analyses, k_off_ is the most influential parameter, followed by k_on_ and dose. Molecular weight and valency overlap with or are below the dummy parameter and thus can be deemed non-influential. There are also additional signs of time-dependent variation, with the influence of k_on_ noticeably peaking at around 4 hours for the elapid.

These results indicate the primary influence of k_on_ on the AUC-OT treatment metric. Dose and k_off_ also have a significant influential effect on treatment outcome, with k_off_ having a particularly strong effect on the dynamics of poorly-performing antivenoms. Molecular weight has a much more limited influence, which is mostly seen on the dynamics of therapeutically active antivenoms. Valency has a non-significant impact across all GSAs. The overall patterns of sensitivity are largely similar for the two venom types. An RBD-FAST GSA was performed to test the reproducibility of this analysis, and produced essentially the same rankings (ESM Section 11). To test for potential parameter interactions, we additionally generated scatter plots linking parameter pairs to AUC-OT output values. These plots indicated a noticeable interaction between k_on_ and k_off_, and a slight interaction between dose and affinity (ESM Section 13).

### 3.10 Properties and effects of antivenom affinity across treatment time delays and scales

We looked more closely at the impact of k_on_ and k_off_ on treatment outcome. Analysis of the antivenoms resulting in the lowest 1% of AUC-OT scores at different treatment timepoints showed that the k_on_ mode of these scaffolds increased with increasing treatment delays (ESM Section 12). A plot of these k_on_ modes against antivenom administration time is shown in Figure 11a. The trend is the same for both the elapid and the viper venom, however the latter reaches a higher modal plateau. This suggests that antivenoms with a k_on_ of 1×10^4^ – 1×10^5^ M^-1^s^-1^ are preferential for treatment delays of 1 hour, whereas antivenoms with a k_on_ of over 1×10^5^ M^-1^s^-1^ are preferential for treatment delays of over 1 hour.

**Figure 11.**
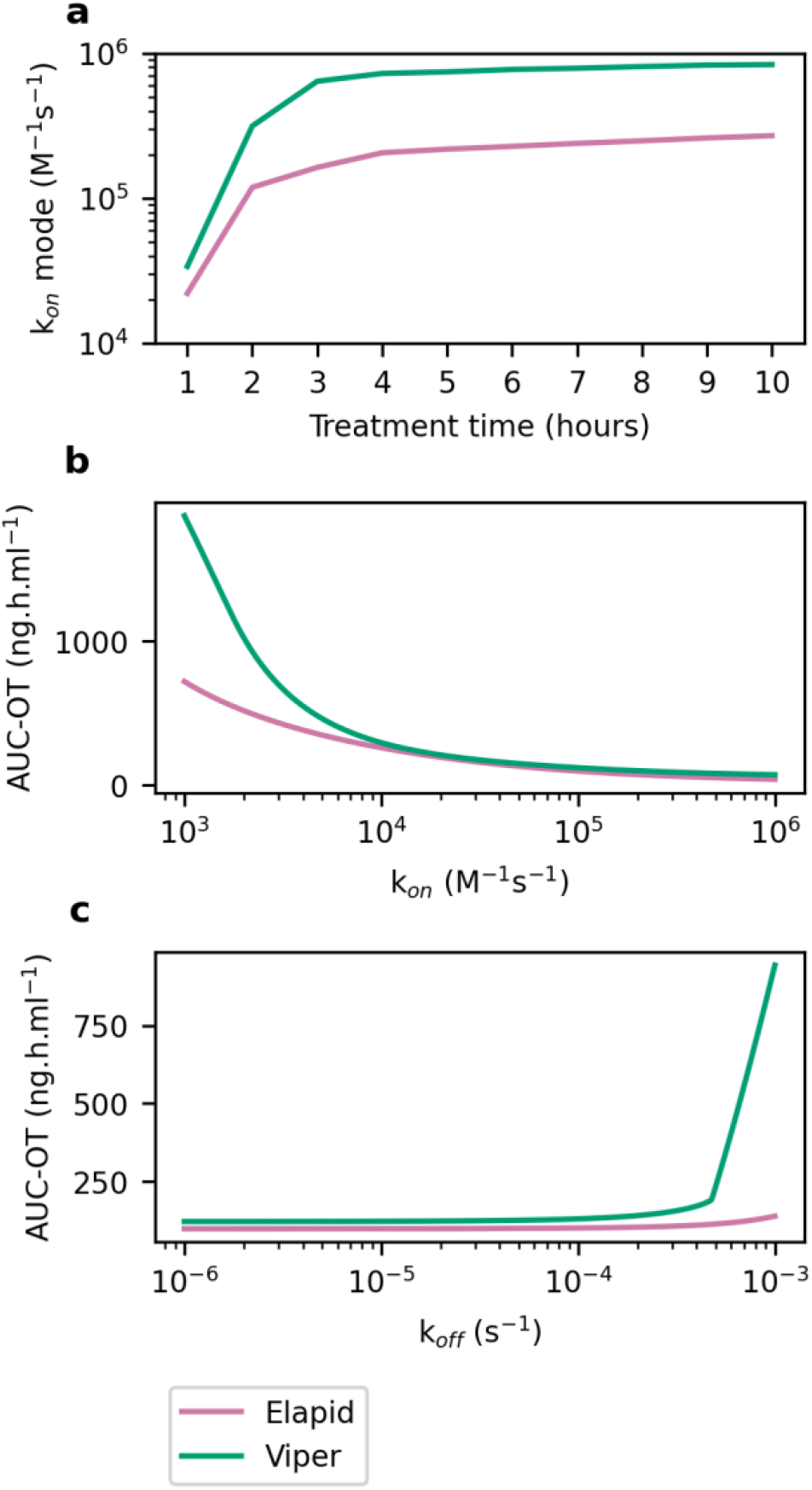
Impact of treatment time on k_on_ mode of effective antivenoms, and LSA of k_on_ and k_off_. Panel a shows that the k_on_ mode of the scaffolds with the lowest 1% AUC-OT scores increases with increasing treatment delays. Panels b and c show the effect of changing k_on_ and k_off_ on AUC-OT. k_on_ variation can be seen to have a greater impact on the AUC-OT range, indicating its greater influence on treatment outcome. In these simulations, 3 mg venom was treated with a 50 kDa monovalent antivenom in a 1:5 venom: antivenom ratio, 3 hours post-bite. For panel b, k_off_ was fixed to 1×10^−5^ s^-1^ and k_on_ was varied between 10^3^ – 10^6^ M^-1^s^-1^. For panel c, k_on_ was fixed to 1×10^5^ M^-1^s^-1^, and k_off_ was varied between 10^−6^ – 10^−3^ s^-1^.

We next performed an LSA of k_on_ and k_off_ on model output, to visualise the effects of varying each parameter individually (Figure 11b-c). These plots show the greater impact of k_on_ on treatment outcome, with k_on_ variation mediating a larger impact on the AUC-OT metric than k_off_. The AUC-OT scores also plateau further along the tested k_on_ range compared to k_off_. This indicates that there are limited benefits of decreasing k_off_ below around 1×10^−4^s^-1^, whereas AUC-OT improvements are still seen with k_on_ increases of up to around 1×10^6^ M^-1^s^-1^.

## 4 Discussion

### 4.1 A computational framework to optimise antivenom design

Computational modelling can be used to improve our understanding of venom-antivenom pharmacodynamics and develop effective treatments. In this study, we applied a computational framework to explore antivenom parameter space and assess the influence of five design features on the treatment of systemic snakebite envenomation. We compared treatment outcomes for a model elapid and viper snakebite to elucidate variations arising from the differing pharmacokinetics of low and high molecular weight venoms, respectively. The model equations are generalised and can in future be applied to additional snake species. We defined AUC-OT as a generalised, continuous metric to quantitatively assess treatment outcome for venoms with different pathologies. The AUC-OT scores resulting from a range of treatment simulations were used to identify the parametric trends associated with effective antivenom treatment. We also undertook a GSA to rank the influence of different antivenom design parameters on treatment outcome. Predictions generated through this modelling approach can inform and be informed by experimental work in a Design-Build-Test-Learn cycle. The parallel testing of different treatment scenarios including time delay and venom type can enable the design of therapeutics for specific or generalised scenarios. These simulation principles could be used to optimise other envenomation treatments, including oligoclonal antivenom cocktails of multiple scaffold types, and combinatorial therapies involving repurposed small molecule enzyme inhibitors (Bulfone et al., 2018; Kini et al., 2018).

### 4.2 Guidelines for effective antivenom design

The results from our model provide several general guidelines for the effective pharmacodynamic design of antivenoms:

1. **There is significant flexibility in the design of effective antivenoms**. Effective antivenoms spanned a wide area of the tested parameter space.
2. **Treatment outcome is primarily mediated by antivenom affinity**. High k_on_ rates (>1×10^5^ M^-1^s^-1^) are required for the most effective treatments. k_on_ has the largest impact on therapeutically active antivenoms and the system as a whole, whereas k_off_ has the largest impact on poorly-neutralising antivenoms. There are limited benefits to reducing k_off_ beyond 1×10^−4^ s^-1^.
3. **Antivenom molecular weight has a minimal direct impact on treatment outcome, however low molecular weight scaffolds offer more flexible design ranges for improved treatment**. Effective antivenoms can be found across a 15 – 150 kDa size range. Low molecular weight scaffolds are increasingly preferable with delayed treatment times.
4. **Low molecular weight scaffolds require higher baseline doses to counteract their shorter half-life, however out-perform high molecular weight scaffolds when dosed sufficiently**. Higher doses generally improve treatment, and bivalency can reduce minimal dose requirements. In the most highly performing treatment grades, low molecular weight scaffolds can tolerate lower doses.
5. **Viper and elapid envenomation systems show the same sensitivity patterns and can be effectively treated by the same types of scaffold**. Delayed treatment imparts more stringent constraints on the parameter bounds of the most effective viper antivenoms, compared to elapid antivenoms.

Taken together, the model suggests that an antivenom that could effectively treat both elapid and viper envenomation at early and delayed treatment times would be small (20 – 30 kDa), high affinity (k_on_ at or above 1×10^5^ M^-1^s^-1^, k_off_ at or below 1×10^−4^ s^-1^), and either monovalent or bivalent. A bivalent antivenom would have lower minimal dosing ratios of at least 1:2 for the elapid, and 1:5 for the viper. It is worth noting that while we defined the most effective scaffolds as those resulting in the lowest 1% AUC-OT scores to easily visualise the underlying trends, a more relaxed threshold would likely suffice. This would expand the design space further, for example by allowing lower dosing ratios. Even with high dosing ratios, the use of low molecular weight scaffolds would likely enable significantly lower absolute mass doses of antivenom. This effect would be compounded by the higher therapeutic efficacy of recombinant antivenoms compared to serum antivenoms, which frequently contain large proportions of non-therapeutic antibodies. Large scaffolds can permit lower dose ratios compared to smaller scaffolds due to their longer half-life, however treatment outcomes are poorer compared to small scaffolds beyond certain dosing ratios. Higher minimal doses of low molecular weight antivenoms are required to compensate for their shortened half-life and see the full pharmacokinetic benefit of their increased tissue distribution.

Monovalent scFvs (27kDa) and bivalent nanobodies (∼30 kDa) are two scaffolds which could potentially meet the design guidelines suggested by our model. scFvs and monovalent nanobodies have already been successfully selected to neutralise a number of snake venom toxins (Fernandes et al., 2021; Laustsen et al., 2018). Bivalent and multivalent nanobody constructs targeting snake and scorpion venom toxins have also been produced (Hmila et al., 2010, 2008; Wade et al., 2022). While scFvs are well-established therapeutic scaffolds, nanobodies have superior thermostability and solubility, lower production costs, and can be more readily conjugated to produce multivalent and multispecific binders (Arbabi-Ghahroudi, 2017; Asaadi et al., 2021; Wang et al., 2022). Nanobodies have naturally low immunogenicity, despite being derived from camelid immunoglobulins, and can be humanised to further improve their safety (Ackaert et al., 2021; Vincke et al., 2009). These properties may make nanobody scaffolds preferable. The scaffold solutions predicted by the model would have to be screened on a case-by-case basis for venom-specific exceptions. For example, nanobodies preferentially bind to clefts and grooves (for example enzyme active sites) and scFvs to flat epitopes, meaning that they may better suit different toxin types (Asaadi et al., 2021; Gilbreth and Koide, 2012). Steric hindrance may also make certain combinations of multivalent binders and toxin ligands unfeasible.

While we were unable to find experimental studies comparing the *in vivo* function of nanobody and scFv antivenoms with conventional scaffolds, there is evidence to support a tolerance of differently sized conventional antivenoms in envenomation treatment. Several studies have described the similar treatment of viper envenomation with Fab, F(ab’)_2_, and IgG scaffolds, in both human and animal studies (Carotenuto et al., 2021; Chaves et al., 2003; Dart and McNally, 2001; Gerardo et al., 2021; León et al., 2001, 1997; Mascarenas et al., 2020). One study also describes the effective treatment of an elapid venom with both IgG and F(ab’)_2_ antivenoms (León et al., 1999). Notably, of the 17 studies we found comparing the effect of different antivenom scaffolds in envenomation treatment, only 2 assessed low molecular weight venoms (from elapid snakes and scorpions). More research on the efficacy of different treatments for elapid envenomation is thus required.

The cost of manufacture is an important consideration in antivenom design due to the prevalence of snakebite in low-income countries. Recombinantly produced IgG antivenoms have been predicted to be cost-competitive with existing serum-based antivenoms, with alternative scaffolds such as nanobodies potentially offering lower costs due to their smaller size and potential for production via microbial fermentation (Jenkins and Laustsen, 2020).

Alternative scaffolds may however be more difficult to clinically approve due to their less well-established history of use in humans. Several obstacles to clinical translation have been reported for nanobody therapeutics (Yang and Shah, 2020). It is also worth noting that production of recombinant polyvalent cocktail antivenoms is anticipated to be more expensive than their monovalent counterparts, and would likely face more hurdles in manufacturing, formulation, clinical trial design, dosing, and regulation (Jenkins and Laustsen, 2020; Larbouret et al., 2021).

### 4.3 Future work and applications of pharmacokinetic modelling in antivenom design

More experimental research is required to characterise and compare the pharmacokinetic properties of different antivenom scaffolds, particularly non-conventional scaffolds. Standardised studies of different antivenoms both in isolation and in envenomation-treatment systems would improve our pharmacokinetic parameter prediction and enable further validation of the model. We were unable to find data on toxicity limits of the different scaffolds in rabbits: knowledge of this would also allow us to refine the dosing ratios tested. To assess the impact of antivenom format in human envenomation treatment, human pharmacokinetic parameters would need to be used. Since this data is difficult to obtain for toxins in humans due to clinical study limitations, parameters for venoms could be obtained through allometrically scaling parameters from animals, or through approximating their value using parameters from similarly-sized proteins. Machine learning approaches are also being used to predict the pharmacokinetic parameters of small molecule drugs (Chou and Lin, 2022; Miljković et al., 2021; Ota and Yamashita, 2022). Similar methods could in future be applied to predict protein pharmacokinetics, with a machine learning model having been recently developed to predict the subcutaneous bioavailability of monoclonal antibodies (Lou and Hageman, 2021)

Tailored studies grading the severity of symptoms at different venom concentrations would also improve the definition of the AUC-OT threshold. Defining a threshold for damage was based on blood concentrations from clinical studies which do not necessarily translate to peripheral compartment thresholds, and which are subject to variation based on the patient’s treatment time delay. Other metrics could be explored in future, including the TOT metric or pathology-specific metrics tracking the levels of specific envenomation markers. Different types of metrics may be more suited to different types of venoms. A mathematical model tracking creatine kinase levels has been defined to describe the onset of myotoxicity following *Pseudechis porphyriacus* envenomation (Sanhajariya et al., 2021). A similar approach could be used to incorporate greater mechanistic detail, which could allow the model to directly interface with experimental antivenom efficacy assays.

Compartmental ODE models have specific advantages in that they are simple and thus computationally efficient, they have limited parameters which reduces error from parameter estimation, and they can describe bulk system dynamics. The assumption of homogenous compartmental mixing however may not accurately describe more granular interactions that occur on the tissue level. Agent-based simulations which track the movement of toxins and antitoxins through space could in future be used to represent the pharmacokinetics in specific tissues and to simulate the onset and treatment of localised tissue necrosis. Local tissue simulations could be interfaced with our systemic model in a multiscale modelling approach, to offer a more complete pathological description of envenomation. Spatialised simulations have been successfully applied to the design and optimisation of cancer-targeting nanoparticles, and similar methodologies could be applied to the neutralisation of venom toxins (Hauert et al., 2013; Stillman et al., 2021, 2020). Spatial models could be validated *in vitro* using organotypic tissue models, which have recently been applied to snakebite envenomation (Ahmadi et al., 2022; Grego et al., 2017). This could help reduce reliance on animal testing within antivenom research, and contribute to the pre-clinical testing of novel antivenom formulations.

Stochastic agent-based models would also be able to capture the microscale dynamics of binding, where the local tissue environment can either hinder or encourage the efficient retention of toxin targets (Dunlap and Cao, 2022). Our model highlights the importance of affinity dynamics, with k_on_ identified as having the greatest influence on treatment outcome in the GSA. The importance of k_on_ over k_off_ is likely due to its ability to not only increase the speed of toxin capture, but also the speed of rebinding following dissociation, which effectively increases occupancy time (Vauquelin, 2016). Interestingly, valency had a non-significant impact across all GSA simulations. We believe that the low influence of the valency parameter within the model may be due to the lack of spatialisation, since both binding sites in a bivalent antivenom are subject to independent saturation dynamics. Incorporation of spatialisation would improve the simulation of localised toxin binding and rebinding, which could increase the influence of the valency parameter.

## 5 Conclusions

Due to the differing pharmacokinetic properties of venoms and antivenoms, the choice of antivenom scaffold may impact the efficacy of envenomation treatment. In this study, we used a computational model of envenomation to assess the impact of antivenom format on treatment outcome. The model is capable of simulating different venoms, and antivenoms of variable sizes, valencies, and affinities. To elucidate the system dynamics underlying envenomation-treatment systems, we conducted a range of simulations to compare different antivenoms in the treatment of two pharmacokinetically distinct venoms. We implemented a global sensitivity analysis and parameter optimisation of antivenom pharmacodynamics to identify the most influential antivenom parameters and their bounds for effective treatment.

These simulations have demonstrated the primary importance of affinity, in particular the on-binding rate, in suppressing toxin concentrations. The molecular weight of the antivenom was found to have a limited direct impact on treatment outcome, however low molecular weight scaffolds appear to offer the most flexibility in the effective treatment of both low and high molecular weight venoms, in early and delayed treatment scenarios. The same antivenom preferences and trends were found for the model elapid and viper envenomation cases, however effective viper antivenoms showed tighter parameter constraints. This computational methodology can be used to inform future antivenom optimisation studies and guide the design and selection of next-generation antivenoms for varied venom types.

## Supporting information

Electronic Supplementary Materials

## 6 Acknowledgements

This work was carried out using the computational facilities of the *Advanced Computing Research Centre*, University of Bristol - http://www.bristol.ac.uk/acrc/.

## Abbreviations

AUC: area under the curve
AUC-OT: Area under the curve, over a threshold
CDF: cumulative distribution function
GSA: Global sensitivity analysis
KDE: Kernel density estimation
KS: Kolmogorov-Smirnov test
LSA: Local sensitivity analysis
TOT: time over threshold

## Statements and Declarations

The authors have no relevant financial or non-financial interests to declare.

## Author Contributions

**Natalie Morris: Conceptualization, Methodology, Software, Formal Analysis, Investigation, Visualization, Writing – Original Draft. Johanna Blee: Conceptualization, Methodology, Writing – Review & Editing, Supervision. Sabine Hauert: Conceptualization, Methodology, Writing – Review & Editing, Supervision**.

