## Supplementary material for "Global parameter optimisation and sensitivity analysis of antivenom pharmacokinetics and pharmacodynamics": Electronic Supplementary Materials

### 1 Calculation of venom molecular weight

We approximated venom molecular weight by calculating a weighted average mass of the venom’s constituent toxins, using proteomic proportions as the weightings. The proteomic composition of *Naja sumatrana* venom was taken from Yap et al. (2014), and the composition of *Cryptelytrops purpureomaculatus* from Zainal Abidin et al. (2016). The toxin families, their proportions, and their molecular weights are listed in Tables S1 and S2. Where ranges for toxin family molecular size were listed in the literature, the median was used to represent the toxin molecular weight. The molecular weight of starred toxins (*) was calculated by multiplication of toxin amino acid number, with the average molecular weight of a single amino acid, 110 Da (Promega, 2022). The SVMP molecular weight in *C. purpureomaculatus* venom was taken as the median across PII and PIII subclasses, which were noted to be the dominant forms in this venom. The BCNP family was not included in the *C. purpureomaculatus* calculation, owing to its small size and low proportion. The weighted average molecular weight of *N. sumatrana* venom was 9.25 kDa, and the weighted average molecular weight of *C. purpureomaculatus* venom was 57.7 kDa.

**Table S1: *N. sumatrana* venom toxin composition and molecular weights**

Abbreviations: CTX = cardiotoxin, PLA_2_ = Phospholipase A_2_, LNTX = long neurotoxin, 3FTX = 3 finger toxin

| **Toxin family** | **Proportion (%)** | **Molecular weight (kDa)** | **Median molecular weight (kDa)** | **Molecular weight reference** |
| --- | --- | --- | --- | --- |
| CTX | 44.2 | 6.5 |  | Kumar et al., 1997 |
| PLA_2_ | 30.6 | 13 - 15 | 14 | Ferraz et al., 2019 |
| LNTX * | 11.5 | 7.3 - 8.1 | 7.7 | Ferraz et al., 2019 |
| Short neurotoxin * | 3.5 | 6.3 - 6.8 | 6.55 | Ferraz et al., 2019 |
| Other 3FTX and minor PLA_2_ | 2.5 | 6.3 - 15 | 10.65 | Upper PLA_2_ bound and lower short neurotoxin bound |

**Table S2: *C. purpureomaculatus* venom toxin composition and molecular weights**

Abbreviations: PLA_2_ = Phospholipase A_2_, SVMP = snake venom metalloproteinase, SNACLEC = snake venom C-type lectin, SVSP = snake venom serine protease, LAAO = L-amino acid oxidase, CRVP = cysteine-rich venom protein, PLB = Phospholipase B, BCNP = Bradykinin potentiating and C-type natriuretic peptide, 5NT = 5’-Nucleotidase, QPCT = Glutaminyl-peptide cyclotransferase, VPDE = venom phosphodiesterase.

| **Toxin family** | **Proportion (%)** | **Molecular weight (kDa)** | **Median molecular weight (kDa)** | **Molecular weight reference** |
| --- | --- | --- | --- | --- |
| SVMP (PII & PIII) | 35 | 30-100 | 65 | Olaoba et al., 2020 |
| SNACLEC | 19 | 27 - 29 | 28 | Morita, 2005 |
| SVSP | 12 | 26 - 67 | 46.5 | Ferraz et al., 2019 |
| LAAO | 10 | 120 – 150 | 135 | Ullah, 2020 |
| PLA_2_ | 8 | 13 - 15 | 14 | Ferraz et al., 2019 |
| CRVP | 6 | 26 |  | Jin et al., 2003 |
| PLB | 2 | 55 |  | Ullah and Masood, 2020 |
| BCNP | 2 | <2 |  |  |
| 5NT | 2 | 120 |  | Trummal et al., 2015 |
| QPCT | 2 | 42 |  | Leonardi et al., 2019 |
| VPDE | 2 | 98 - 140 | 119 | Ullah et al., 2019 |

### 2 Treatment outcome metric definitions

Three quantitative metrics for treatment outcome were defined: the venom area under the curve (AUC), the time that the venom concentration remained over a critical threshold (time over threshold - TOT), and the venom AUC over a threshold (AUC-OT). All metrics were recorded from the point of antivenom treatment to the end of the simulation. A schematic of these metrics applied to a peripheral compartment venom curve is shown in Figure S1.

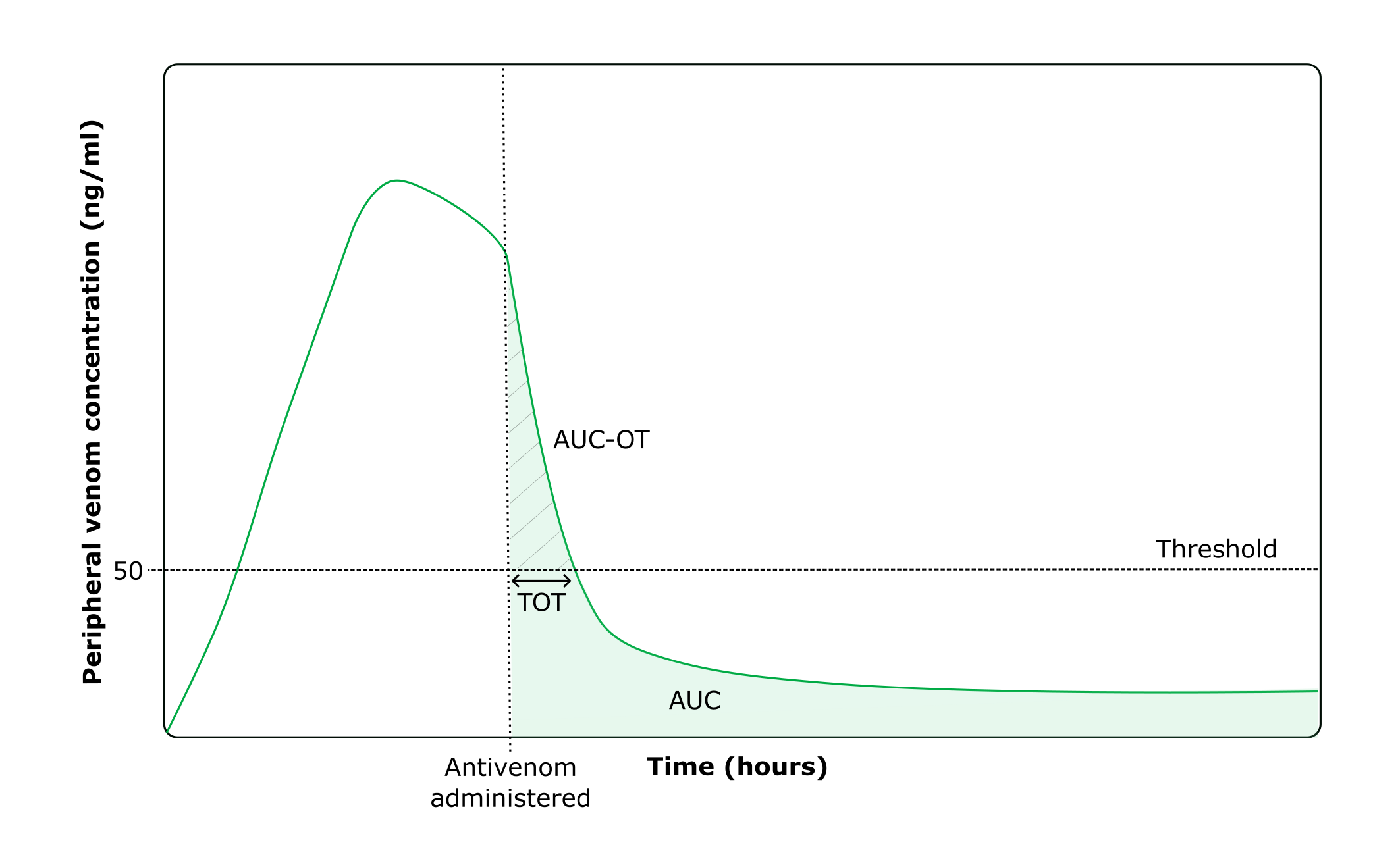

**Figure S1. Schematic of the AUC, AUC-OT, and TOT metric calculations.**

AUC is shown in the green area. AUC-OT is shown in the hatched area. TOT is indicated by the double-headed arrow.

### 3 AUC and AUC-OT Comparison

We compared the AUC and AUC-OT scores in cases studies of the model elapid (*N. sumatrana*) and viper (*C. purpureomaculatus*) envenomation (Fig. S2-S3). All simulations followed a 3 mg intramuscular envenomation, treated with either IgG, F(ab’)_2_, Fab, scFv, or nanobody antivenom in a 1:3 venom: antivenom dose ratio. Antivenom k_on_ was set to 6x10^5^ M^-1^s^-1^, and k_off_ to 1x10^-3^ s^-1^, giving a K_D_ of 1.67 nM. Simulations were repeated for treatment times ranging from 1 to 10 hours post-envenomation.

Fig. S2 shows the central compartment AUC scores, with AUC-OT scores overlain as white hatch marks. Provided the antivenom is of a sufficient dose, the entire AUC score occurs below the threshold leading to an AUC-OT of zero. This is due to the assumptions of homogenous mixing and of an instantaneous antivenom bolus dose. This makes the central compartment AUC-OT metric non-informative in all but the poorest-performing use cases, preventing the effective comparison of scaffolds. We subsequently decided to not use the central compartment AUC-OT.

Fig. S3 shows the peripheral compartment AUC and AUC-OT comparisons. Across all simulations and antivenom types, the majority of the peripheral compartment AUC score occurs below the critical threshold of 50 ng/ml. All of the peripheral compartment simulations result in an AUC-OT score due to the time-delayed absorption of antivenom to the peripheral compartment. This makes this metric more informative for antivenom comparison.

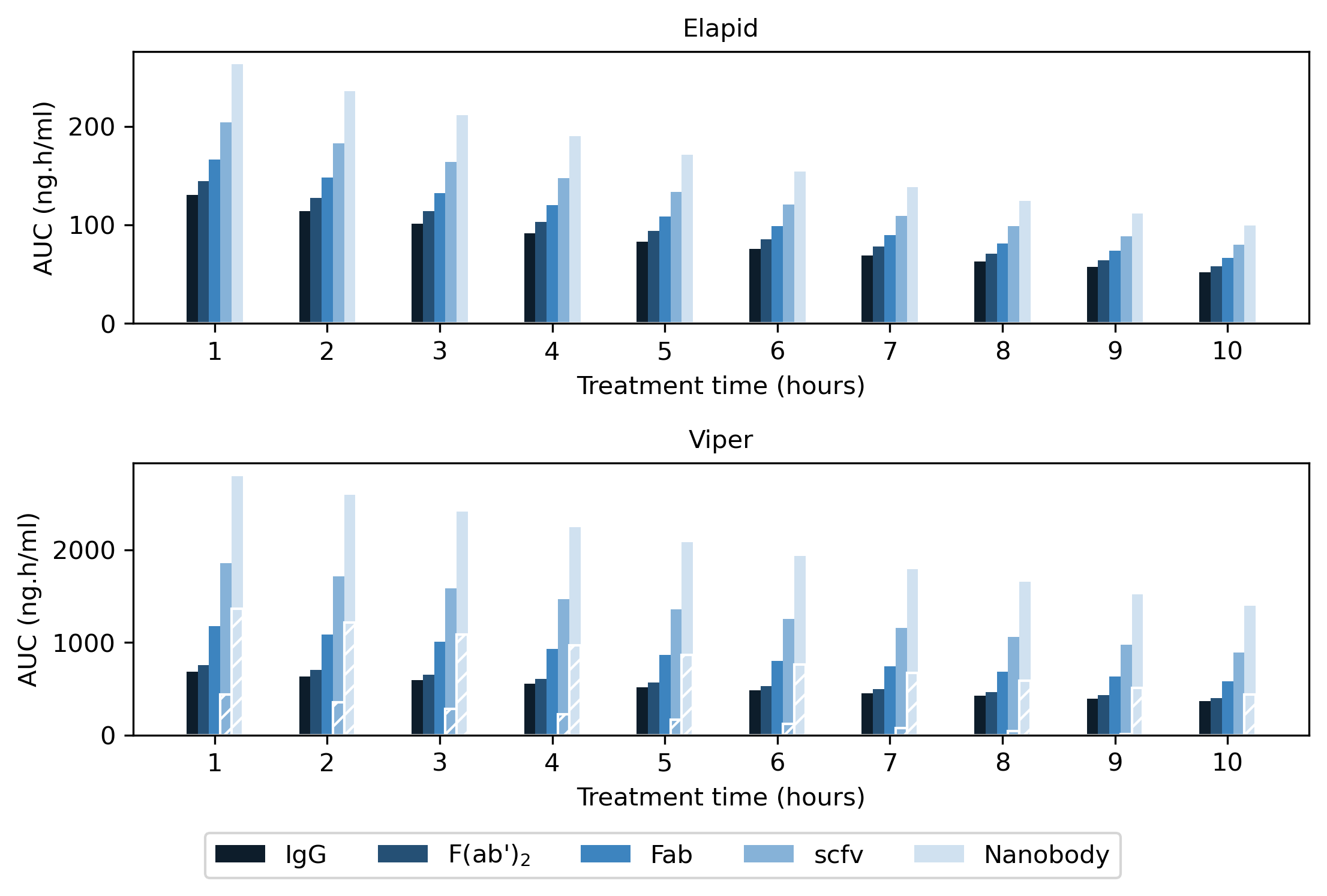

**Figure S2. Central compartment AUC and AUC-OT scores following elapid and viper envenomation.**

Solid bars show the AUC score post treatment, with overlain white hatch marks showing the corresponding AUC-OT score.

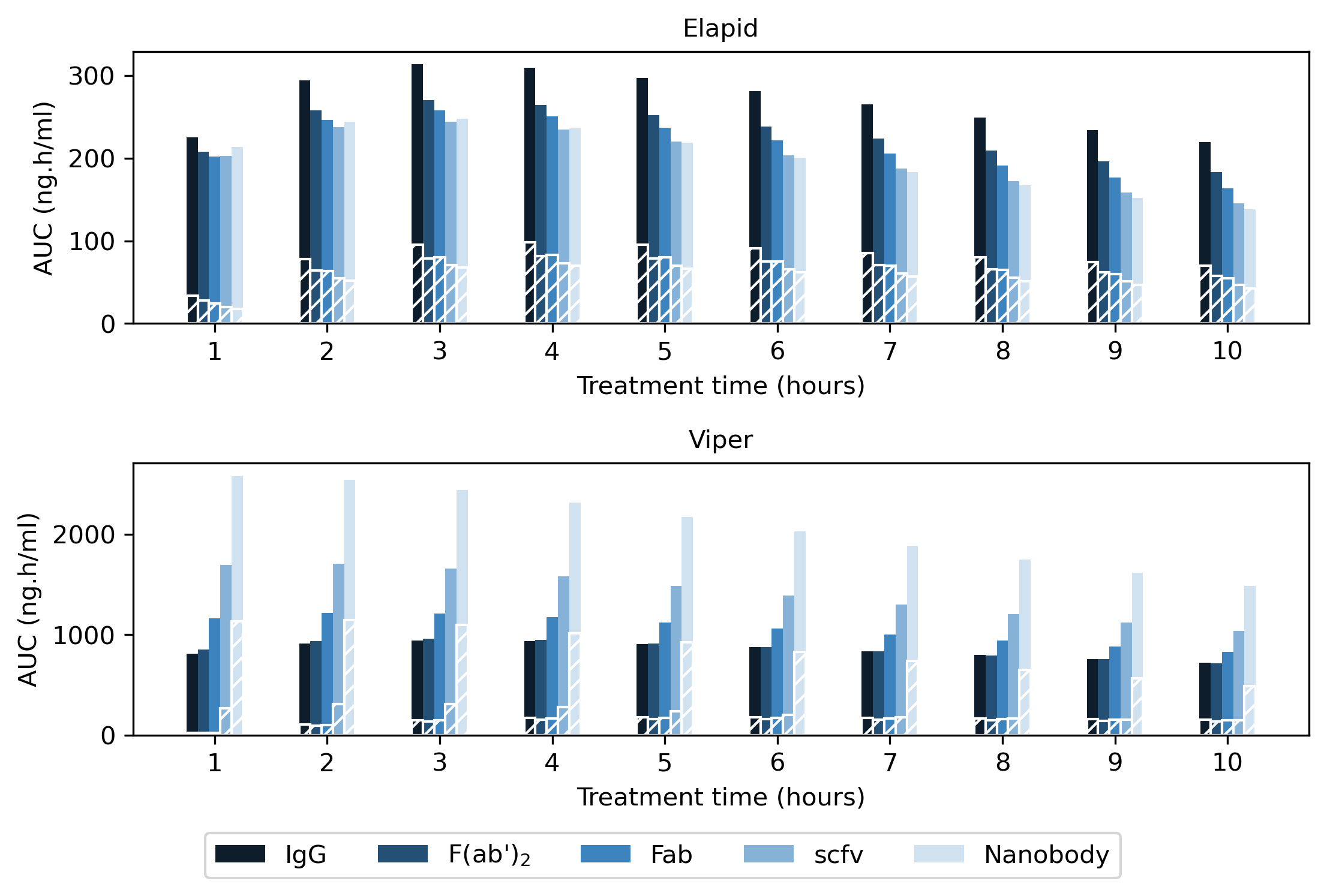

**Figure S3. Peripheral Compartment AUC and AUC-OT scores following elapid and viper envenomation.**

Solid bars show the AUC score post treatment, and with the overlain white hatch marks showing the corresponding AUC-OT score.

### 4 Variable antivenom dosing simulations

We compared the AUC-OT scores following treatment with different antivenoms, dosing ratios, and treatment times. All simulations were performed using a 3 mg intramuscular venom dose, using antivenoms with k_on_ = 6x10^5^ M^-1^ s^-1^ and k_off_ = 1x10^-3^ s^-1^. Fig. S4 shows the elapid simulations, and Fig. S5 shows the viper simulations. Across all simulations, when venom: antivenom dosing ratios are low, smaller scaffolds generally perform more poorly relative to larger scaffolds. Increasing the antivenom dose improves the performance of small scaffolds. At sufficient antivenom doses, there are significant improvements in AUC-OT scores from the use of smaller scaffolds owing to their rapid diffusion into the tissue. The elapid envenomation case is more receptive to treatment with low molecular weight antivenoms, with dosing ratios as low as 1:2 leading to nanobodies outperforming all other scaffolds. The viper envenomation case however requires a higher 1:7 dose for nanobodies to become the preferred scaffold. This is likely due to the longer half-life of viper venom. Equivalent simulations using the TOT metric produced the same trends (data not shown).

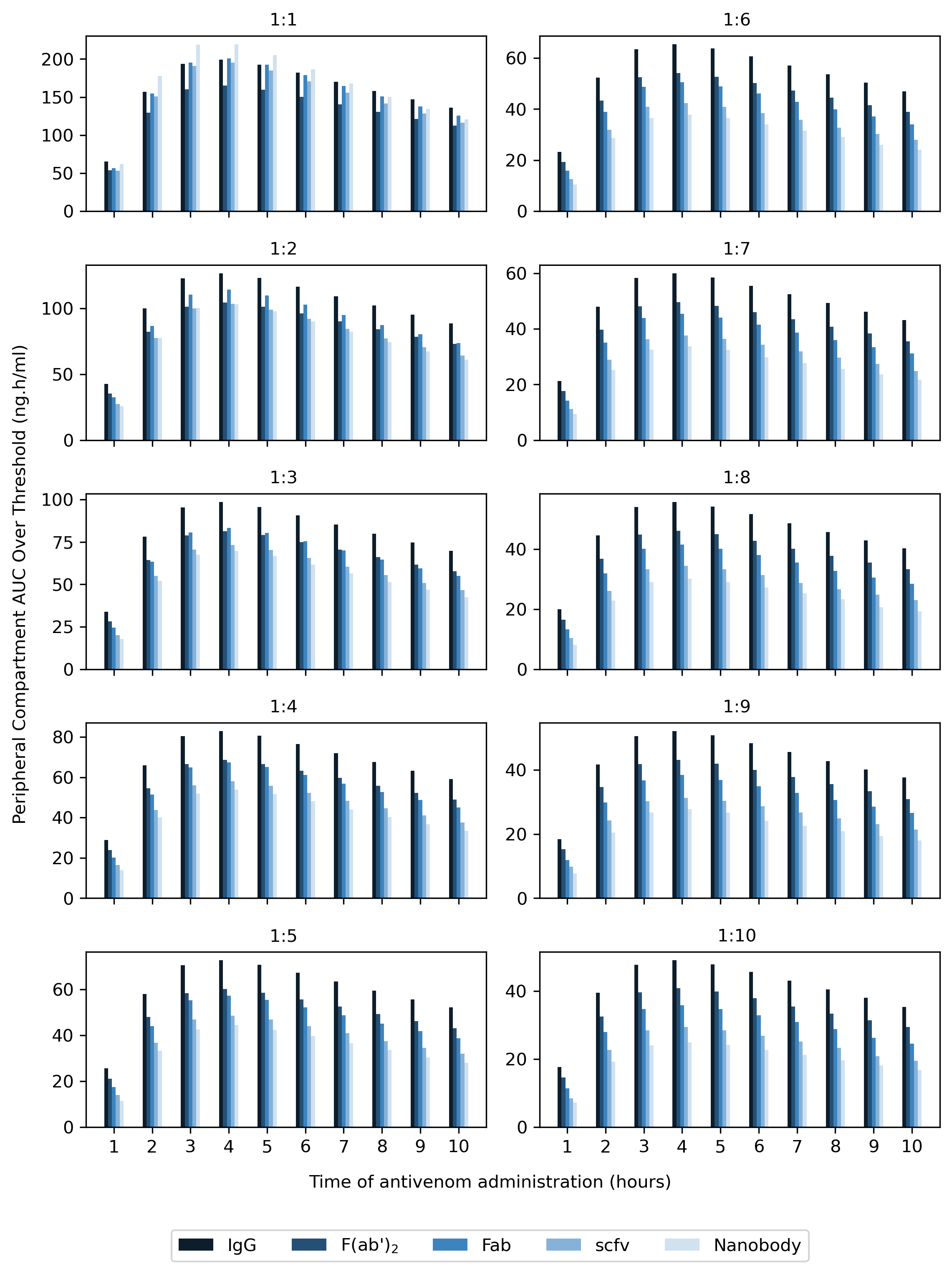

**Figure S4. Elapid envenomation AUC-OT scores following antivenom treatment, at dosing ratios of 1:1 – 1:10 venom: antivenom**

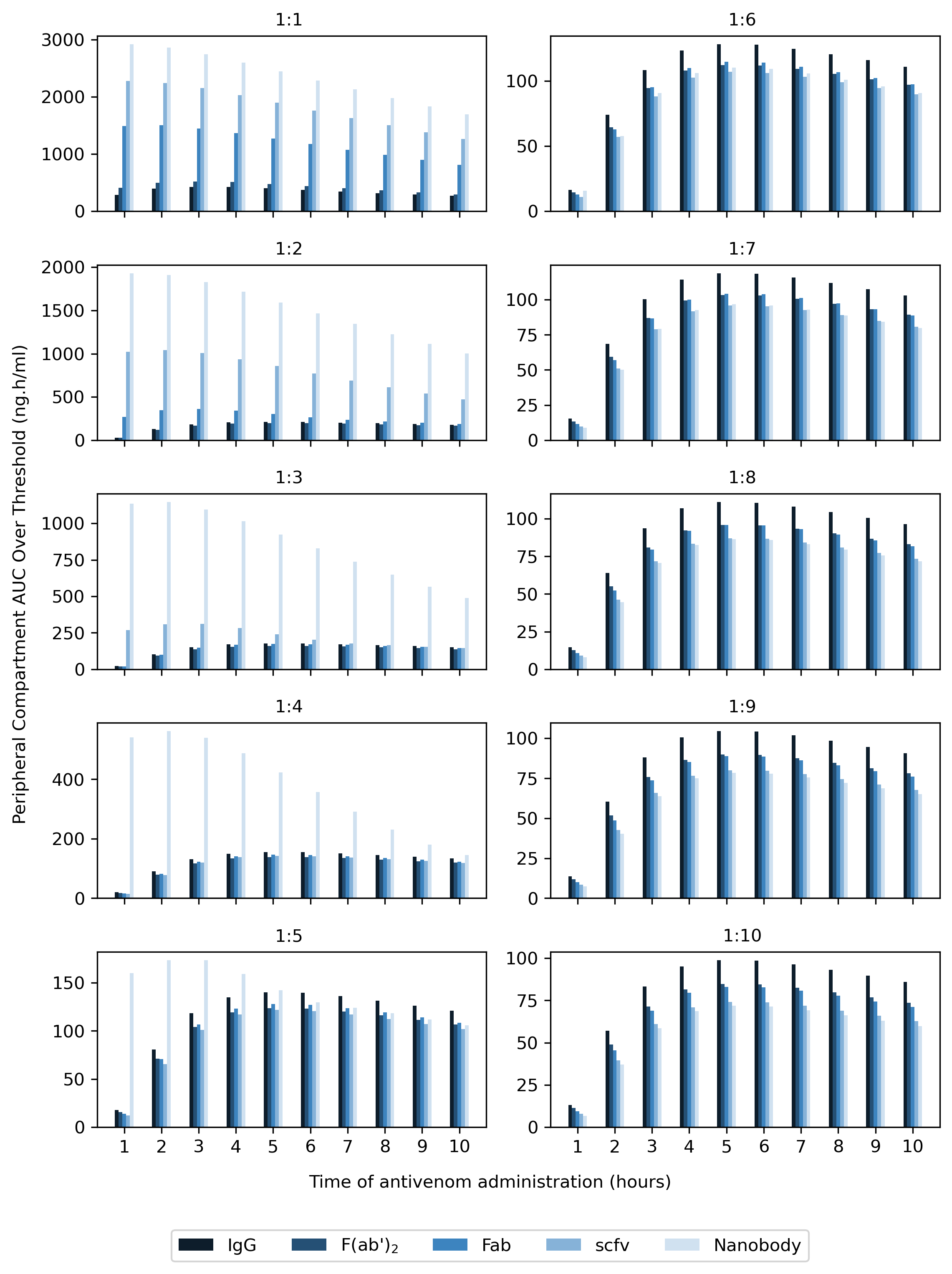

**Figure S5. Viper envenomation AUC-OT scores following antivenom treatment, at dosing ratios of 1:1 – 1:10 venom: antivenom**

### 5 Central compartment AUC analysis

We compared central compartment AUC scores across varying venoms, antivenoms, and treatment times, using the same simulation conditions as ESM Section 4. As Fig. S6 shows, larger scaffolds always lead to lower venom AUC scores in the blood due to their longer half-life. This is true at minimal doses of 1:1 venom: antivenom, and at extreme saturating doses of 1:50.

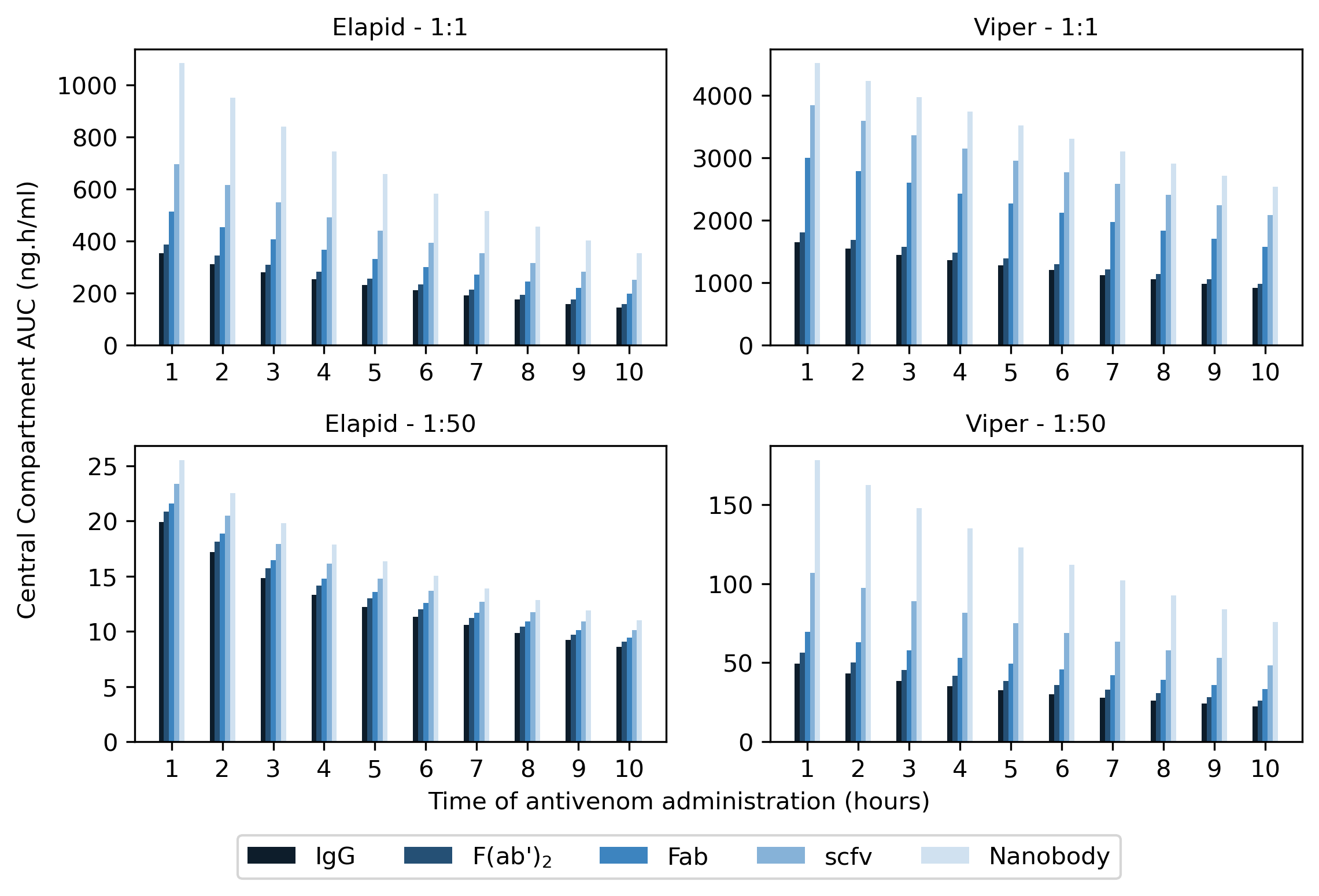

**Figure S6. Central compartment AUC scores following treatment of elapid or viper envenomation with different antivenoms at varying dosing levels.**

k_on_ = 6x10^5^ M^-1^ s^-1^ and k_off_ = 1x10^-3^ s^-1^. Treatment was simulated at various treatment times using dosing ratios of either 1:1 or 1:50 venom: antivenom.

### 6 Theoretical antivenom parameter sample bounds

In setting the upper and lower bounds of our theoretical antivenom parameter set, we aimed to choose physically feasible values. We used an antivenom molecular size range of 15 to 150 kDa and a valency toggle of monovalent or bivalent toxin binding, to allow simulation of a range of antibodies with different valency configurations. It is worth noting that due to the estimation of antivenom k_10_ based on molecular size, antivenoms of 150 kDa in our dataset are simulated with a faster elimination half-life than natural IgGs, whose half-life is extended due to the FcRn recycling system (Ober et al., 2004). While we previously predicted the extended half-life of IgGs to be 84.9 hours, in the theoretical antivenom set 150 kDa scaffolds are predicted to have a half-life of 51.9 hours (Morris et al., 2022).

We used a dosing ratio range of 1:1 to 1:10 venom: antivenom particles. Previous studies in mice have treated low molecular weight dendrotoxins with human IgGs in ratios of up to 1:6 (Laustsen et al., 2018). We opted to increase the maximum dosing limit to examine the effects of highly-saturating antivenom doses. We additionally anticipated that smaller scaffolds may need higher dosing ratios to counteract their shorter half-life. Assuming a 2 kg rabbit size and treatment using the largest antivenom at 150 kDa at a 1:10 venom: antivenom ratio, treatment of 1.5 mg/kg elapid venom leads to a maximum antivenom dose of 105 mg/kg, and treatment of 1.5 mg/kg viper venom leads to an antivenom dose of 16.4 mg. In real life, this may be limited by host immunogenic responses. Doses of 138 mg/kg equine IgG antivenom have previously been applied to rabbits to investigate supportive therapies for adverse antivenom effects (Herrera et al., 2017). Antibody humanisation however could however be applied to recombinant antivenom development to reduce the risk of adverse immune response, which may facilitate future treatment with higher antivenom doses (Harding et al., 2010; Lu et al., 2020). More data would be required to determine the upper dosing limits of different types of antivenom in rabbits.

We used k_on_ bounds of 1x10^3^ – 1x10^6^ M^-1^s^-1^, where 1x10^6^ is the upper limit of naturally occurring antibodies and is controlled by the maximal diffusion coefficients of the antibody and antigen (Foote and Eisen, 1995). k_on_ rates exceeding this ceiling have been found and can result from electrostatic forces between the antibody and antigen (Rathanaswami et al., 2005). We used k_off_ bounds of 1x10^-6^ – 1x10^-3^ s^-1^. While the limit for k_off_ in natural immunity is 1x10^-4^ s^-1^, *in vitro* affinity maturation processes can lead to k_off_ values below this (Foote and Eisen, 1995). The k_on_ and k_off_ parameter bounds resulted in an overall K_D_ range of 10^-12^ - 10^-6^ M, spanning the high micromolar to low picomolar range (from sub-optimal to very high affinity) (Rathanaswami et al., 2005). The weakest affinity combination of k_on_ = 1x10^3^ and k_off_ = 1x10^-3^ s^-1^ generally leads to limited therapeutic activity for elapid, and poor therapeutic activity for viper (data not shown). This affinity combination was thus taken as the lowest boundary, to avoid simulation of large numbers of ineffective antivenoms.

### 7 Output sample space

Figure S7 shows the AUC-OT output distributions of the envenomation-treatment simulations for each snake, at each antivenom administration timepoint. Each graph comprises data from 200,000 simulations. The output distributions across all simulations are heavily right-skewed, indicating that the majority of antivenoms lead to low AUC-OT scores and thus good neutralisation of circulating venom. With delayed treatment times, the distributions become less skewed as the maximum AUC-OT decreases and the parameter ranges conducive to effective treatment narrow.

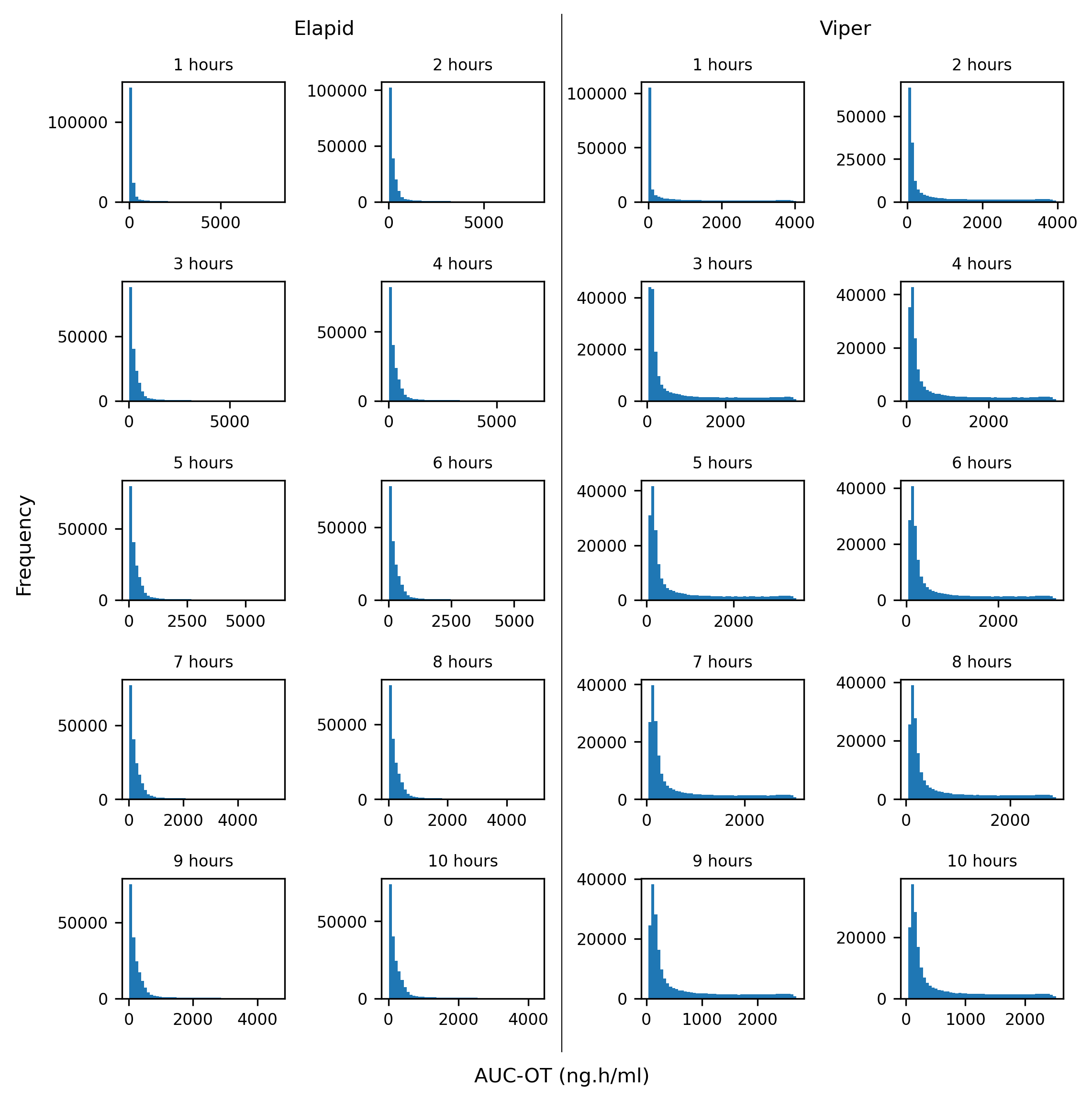

**Figure S7. AUC-OT output distributions for model elapid and viper simulations, at each antivenom administration time.**

### 8 Properties of effective antivenoms at early and late treatment

Histograms were used to identify the theoretical antivenoms which resulted in the lowest 1% AUC-OT scores at each timepoint. The parameters of these scaffolds are shown in pairplots for treatment times of 1 and 6 hours (Figs. S8-11). Across both snake species, increasing treatment time incurs a preference for low molecular weight scaffolds and higher doses. Delayed treatment also incurs a preference for bivalent antivenoms, which is most noticeable for the viper envenomation. Between 1 and 6 hours, the mode of the k_on_ distribution noticeably increases, indicating that lower k_on_ values may be preferable for early antivenom administration. The k_off_ distribution across all figures is relatively static, with a mode of close to 1x10^-5^ s^-1^ apparent in all simulations.

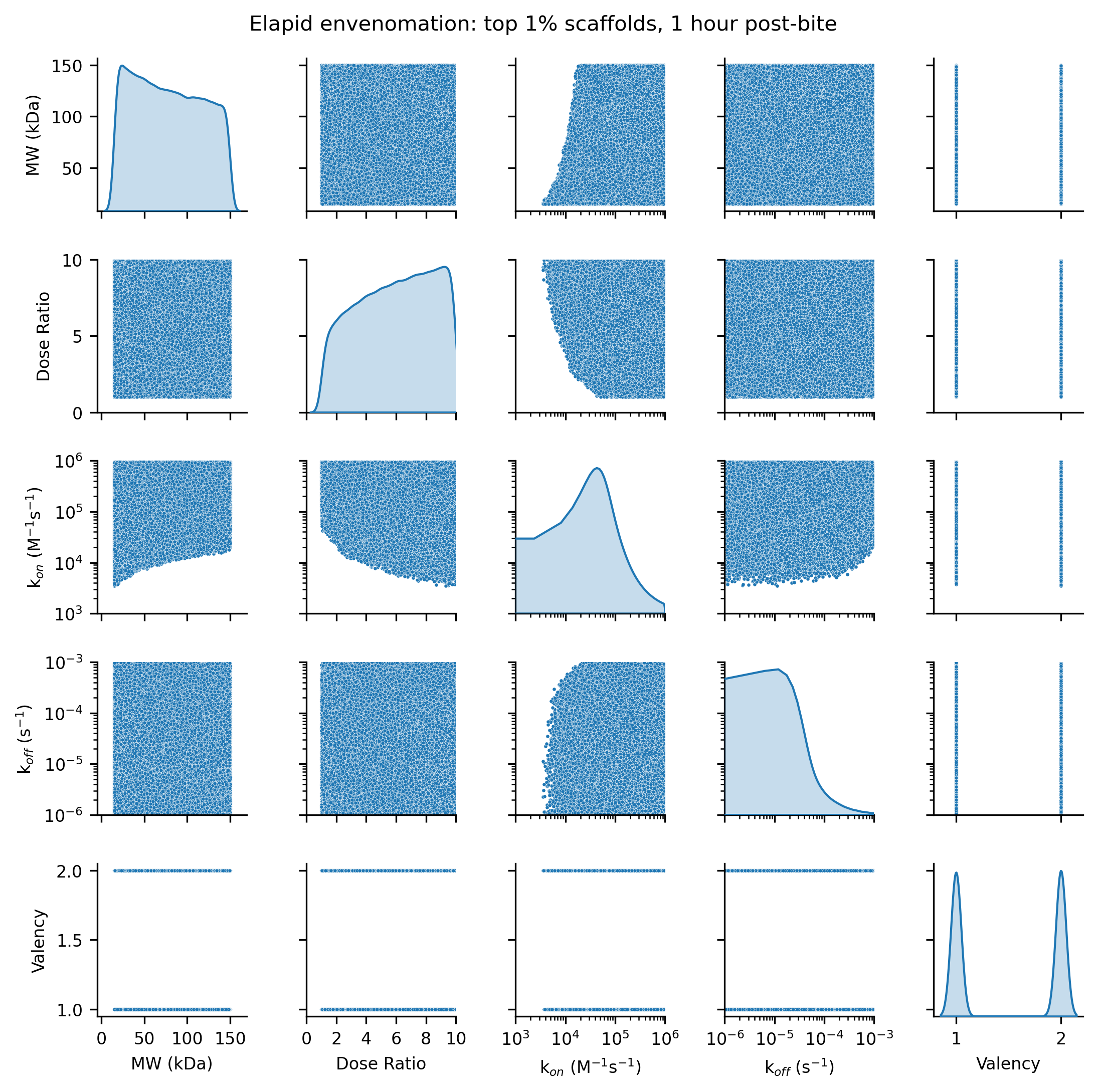

**Figure S8. Pairplot of antivenom parameters resulting in the lowest 1% AUC-OT scores for elapid envenomation treatment, 1 hour post-bite.**

Scatter plots show distributions of paired parameters. Kernel density estimation (KDE) plots along the diagonal show the distributions of individual parameters.

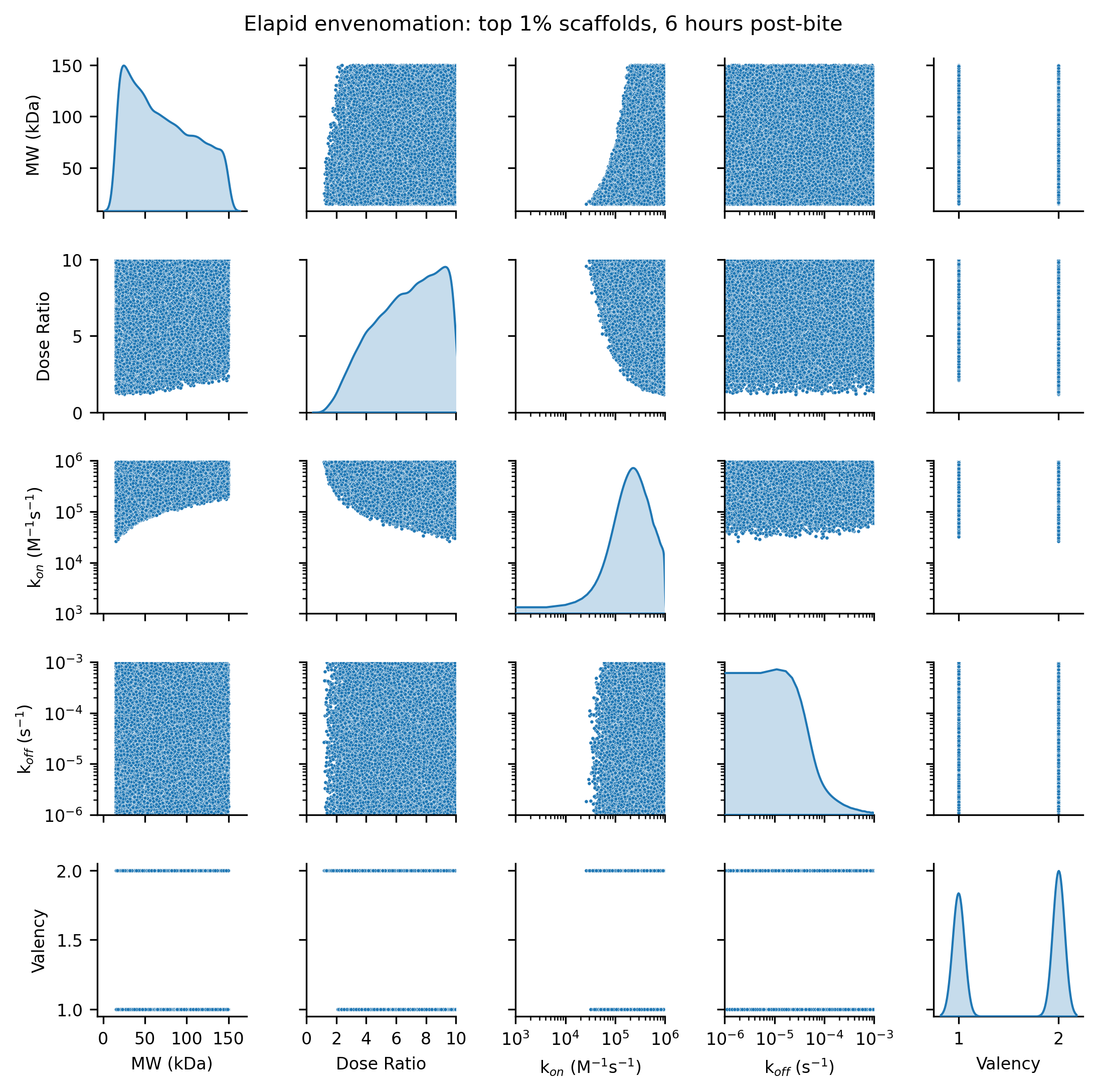

**Figure S9. Pairplot of antivenom parameters resulting in the lowest 1% AUC-OT scores for elapid envenomation treatment, 6 hours post-bite.**

Scatter plots show distributions of paired parameters. KDE plots along the diagonal show the distributions of individual parameters

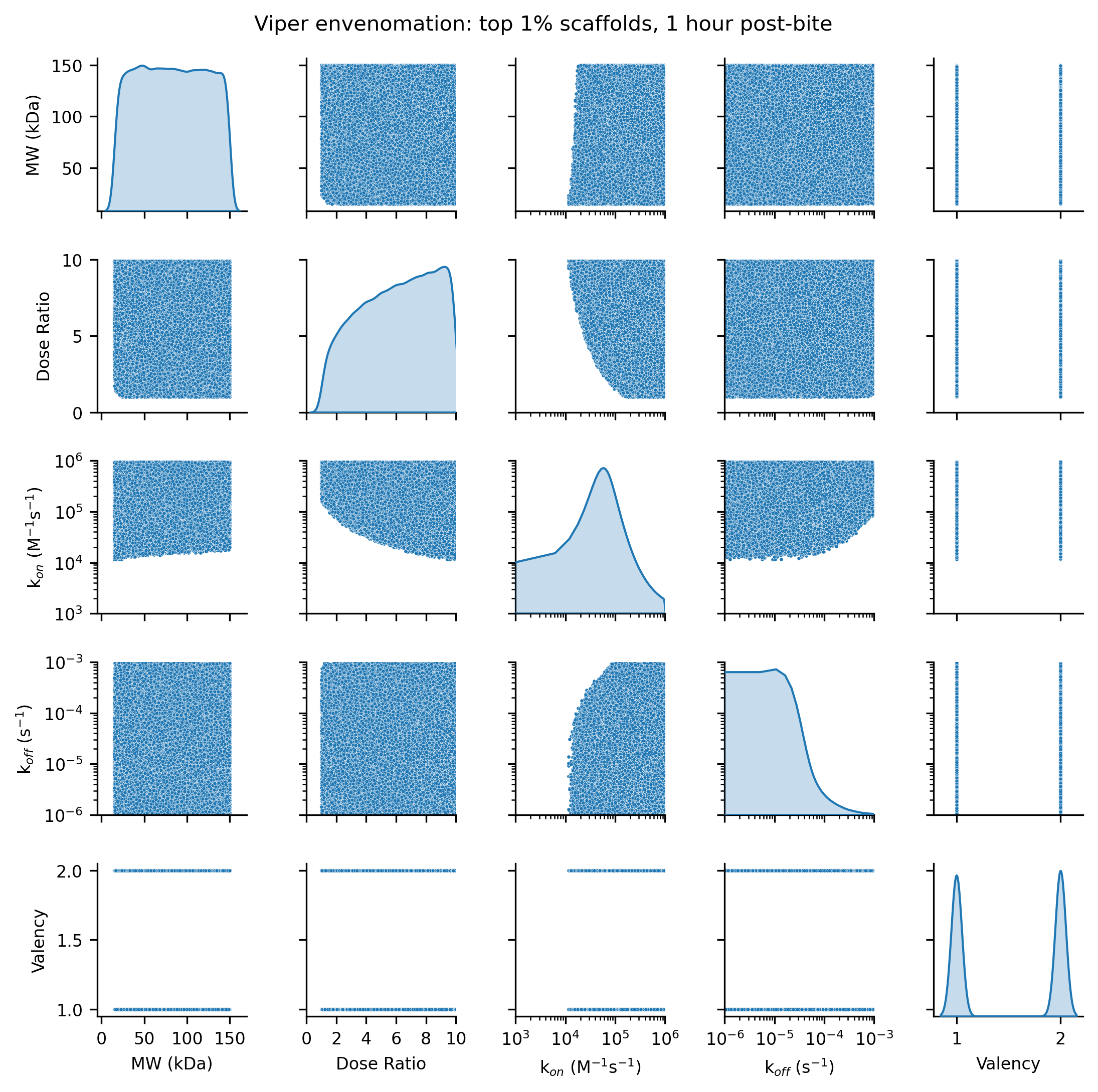

**Figure S10. Pairplot of antivenom parameters resulting in the lowest 1% AUC-OT scores for viper envenomation treatment, 1 hour post-bite.**

Scatter plots show distributions of paired parameters. KDE plots along the diagonal show the distributions of individual parameters.

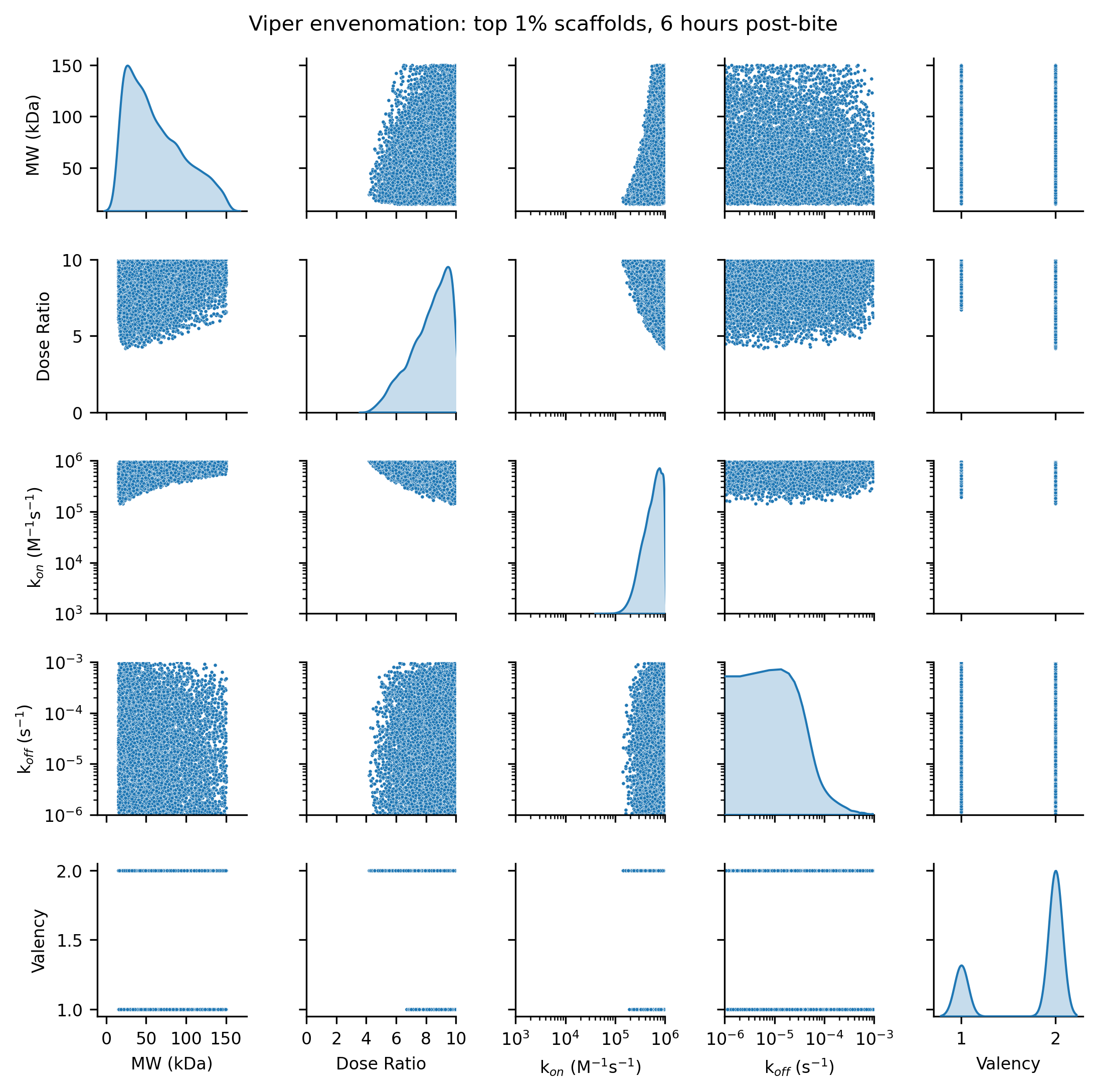

**Figure S11. Pairplot of antivenom parameters resulting in the lowest 1% AUC-OT scores for viper envenomation treatment, 6 hours post-bite.**

Scatter plots show distributions of paired parameters. KDE plots along the diagonal show the distributions of individual parameters.

We next plotted the parameter combinations resulting in the lowest 1% AUC-OT scores at treatment times of 1, 3, and 6 hours (Figure S12)*.* For both snakes, delayed treatment times lead to a constriction in the area of parameter space and a preference for high dose, high affinity, and low molecular weight antivenoms. With greater treatment delays, a greater amount of venom is distributed within the peripheral compartment. This is more rapidly cleared by antivenoms with these characteristics. The shift in parameter space more apparent for the viper envenomation case, with the elapid remaining tolerant of a wider range of scaffolds.

We additionally generated KDE plots of the top 1% performing scaffolds at treatment times of 1 and 6 hours, to visualise the density of antivenoms across parameter space (Figures S13-14). These plots show that in the early 1 hour treatment case, a wide area of area of parameter space is attributable to the top 1% performing scaffolds. While at a 1 hour treatment time, there is a clear preference for low molecular weight scaffolds in the elapid envenomation, there is no strong molecular weight preference for viper envenomation. The changing patterns of density underscore that delayed treatment imparts a preference for low molecular weight scaffolds with high affinity and high dose. The KDE plots also show movement in the k_on_ mode with increasing treatment delay times. For both snakes, at 1 hour the k_on_ mode lies between 1x10^4^ – 1x10^5^ M^-1^s^-1^, whereas at 6 hours post-treatment the mode lies between 1x10^5^ – 1x10^6^ M^-1^s^-1^.

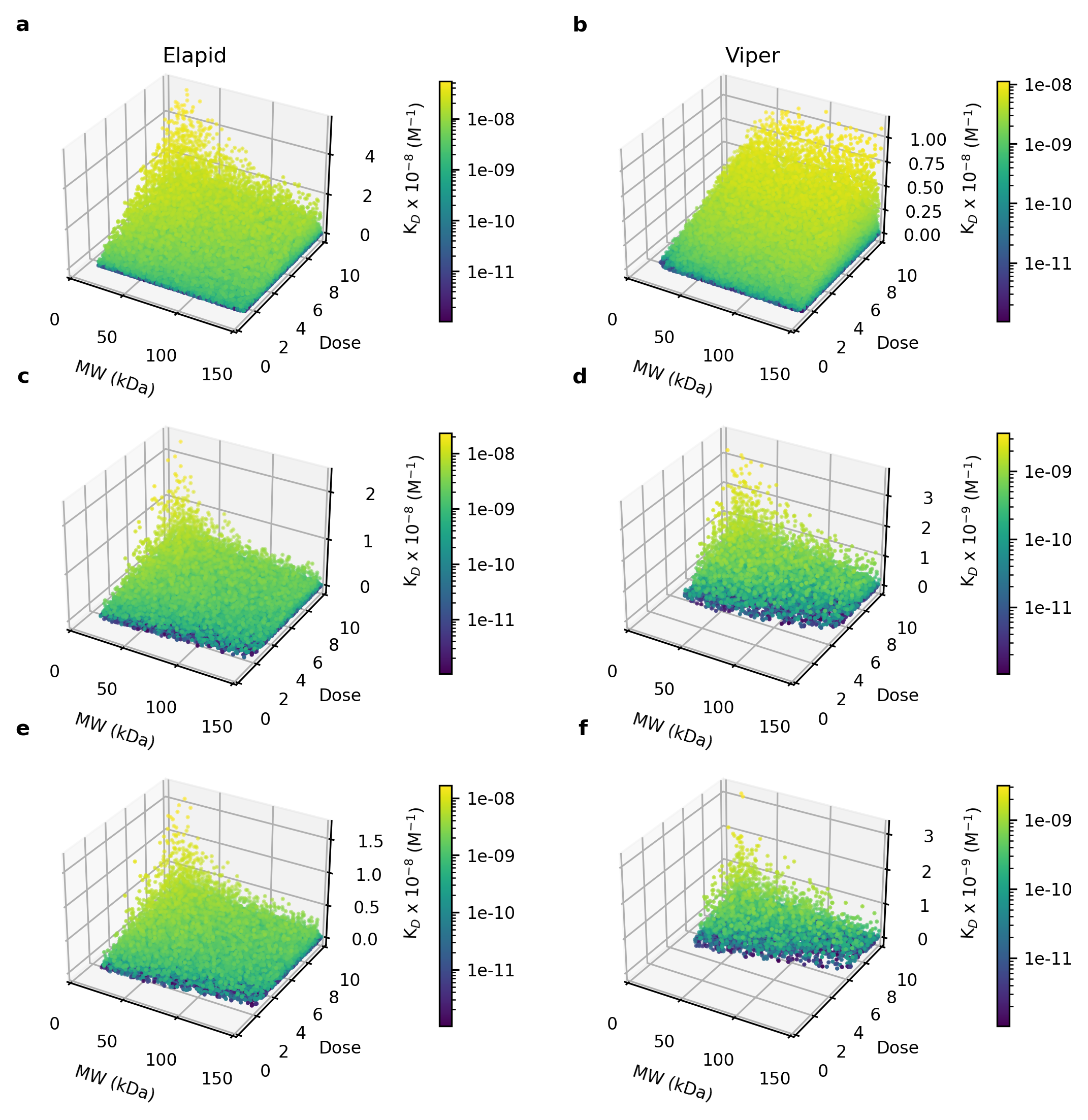

**Figure S12. Antivenom scaffolds leading to lowest 1% AUC-OT scores, at treatment times of 1, 3, and 6 hours post-bite.**

Panels on the left show the elapid results. Panels on the right show the viper results. Panels a-b show the results of treatment at 1 hour, panels c-d show the results of treatment at 3 hours, panels e-f show the results of treatment at 6 hours. The colour bar corresponds to antivenom K_D_ (K_D_ = k_off_/k_on_).

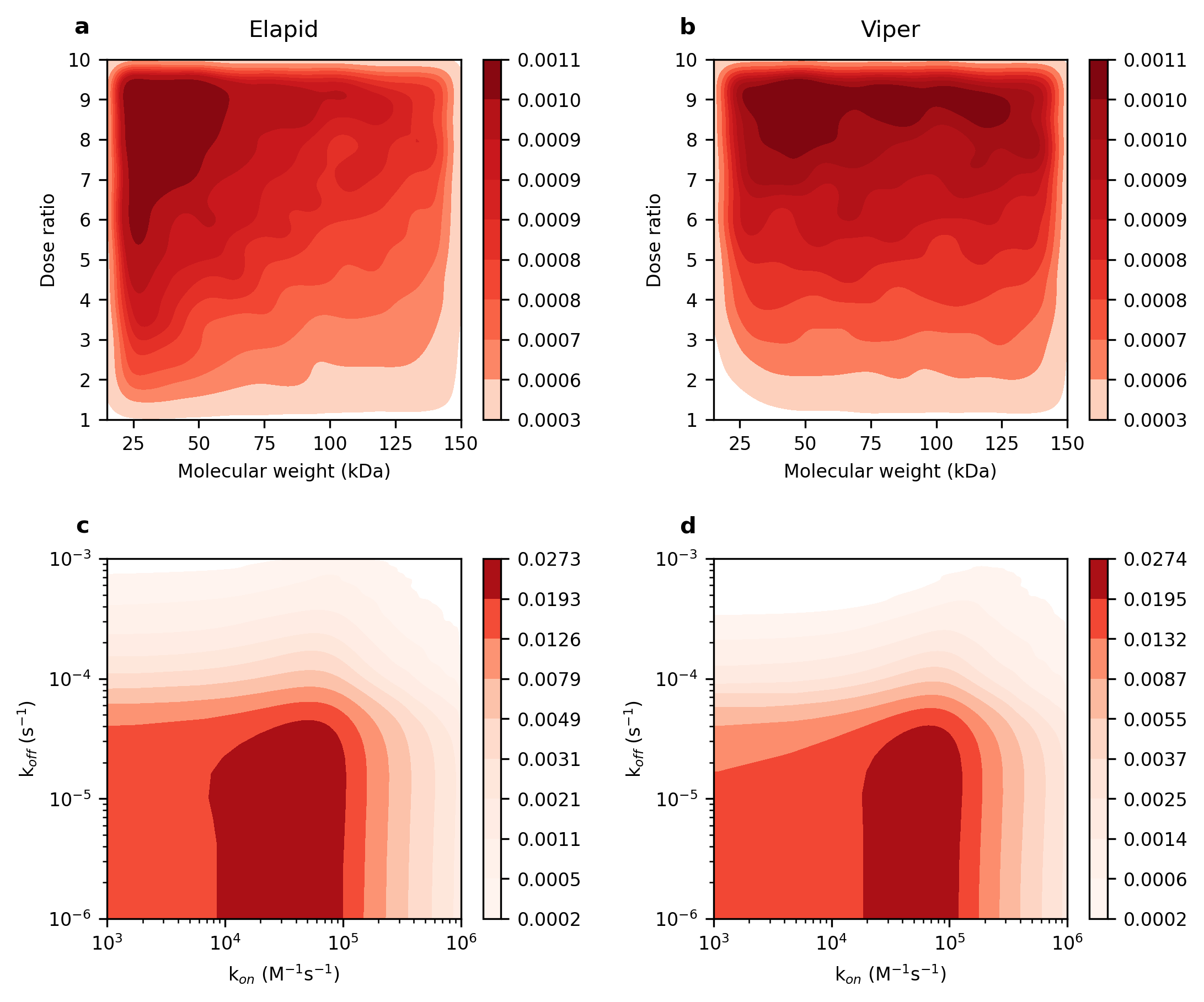

**Figure S13. KDE of top 1% performing scaffolds when antivenom is administered 1 hour post-bite.**

Colour bar corresponds to the KDE. Panels on the left show elapid antivenom parameter space, whereas panels on the right show viper antivenom parameter space.

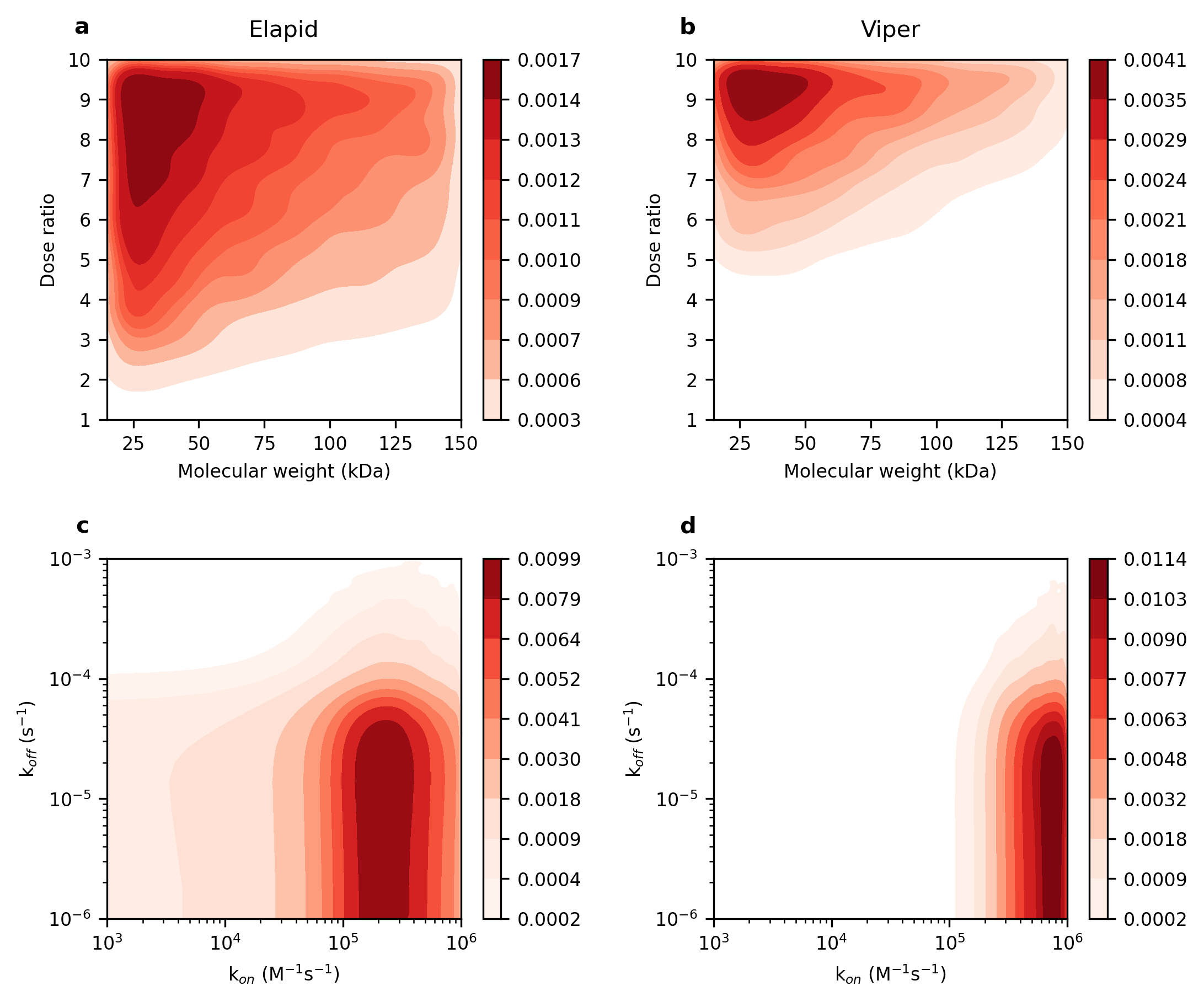

**Figure S14. KDE of top 1% performing scaffolds when antivenom is administered 6 hours post-bite**

Colour bar corresponds to the KDE. Panels on the left show elapid antivenom parameter space, whereas panels on the right show viper antivenom parameter space.

### 9 PAWN Convergence plots

We utilised the PAWN global sensitivity analysis (GSA) method, established by Pianosi and Wagener (2018, 2015). This method compares the cumulative distribution functions (CDFs) of a model’s output when a specific parameter is fixed (in a conditional CDF) with when all parameters are varied (in an unconditional CDF). The difference between CDFs is calculated using the Kolmogorov-Smirnov (KS) test. The KS value represents the PAWN sensitivity index. The more sensitive an output is to a parameter, the greater the difference between the CDFs and the larger the KS value.

We conducted a convergence analysis to ensure a sufficient subsample size for the bootstrapped PAWN. In this, we calculated the median KS over differently sized subsamples of the 200,000 simulations performed at a treatment time of 3 hours. The sample sizes (N) tested were: 500, 1000, 2500, 5000, 7,500, 10,000, 15,000, 20,000, 50,000, 75,000, 100,000, and 200,000. We additionally tested four values of the tuning parameter, n. This designates the number of intervals across which a parameter’s values are allocated, for calculation of the conditional CDFs. It subsequently determines the number of conditional CDFs that are generated. The conditional CDFs are compared to the unconditional CDF, which is generated from the total subsample N and considers simultaneous variation in all parameters. The KS test is then performed to compare each of the n conditional CDFs with the unconditional CDF, and the median KS of these tests is calculated (Noacco et al., 2019; Pianosi and Wagener, 2018).

The resulting convergence plot is shown overleaf (Fig. S15). Comparing plots, changing the tuning parameter n does not have a significant impact on the rankings of parameters. The tuning parameter was subsequently set to 10 for the bootstrapped analysis. For both snakes, a sample size of 7,500 gives a stable ranking that is consistent with larger subsample sizes. For the final analysis, we opted to use 100 bootstrapped resamples of 7,500, which gives very small 95% confidence intervals.

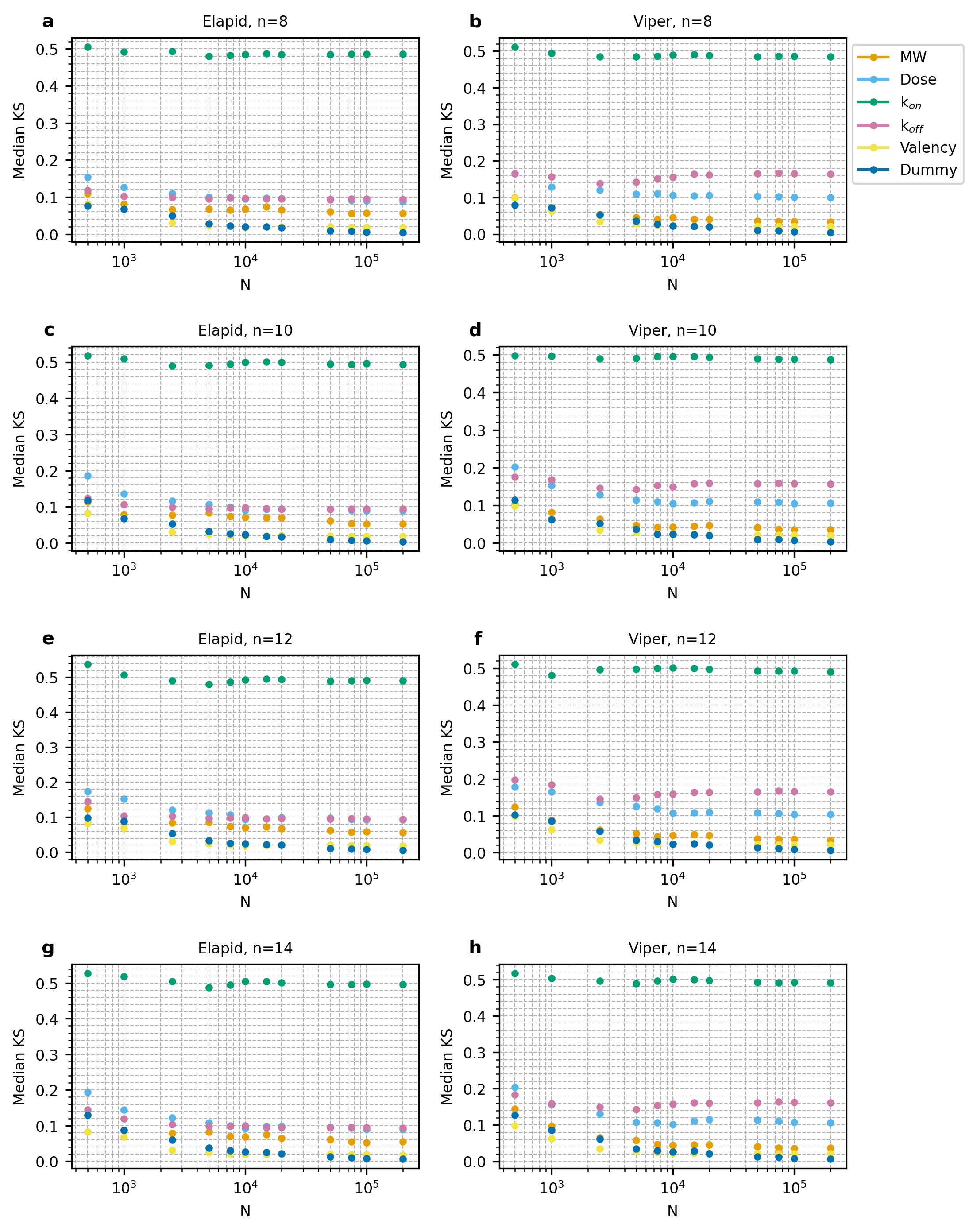

**Figure S15. PAWN GSA convergence plot.**

N = sample size for PAWN index calculation, n = number of conditioning points.

### 10 Effect of uncertain pharmacokinetic parameter prediction on model output

We examined the impact of uncertainty in our pharmacokinetic parameter predictions on the characteristics of effective antivenoms. To do this, we repeated the globally varying antivenom simulations while applying random 2-fold variation to the predicted k_10_, k_12_, and k_21_ values of antivenom and neutralised venom. In these simulations, random 2-fold scaling factors were selected from a uniform distribution, and applied individually to the k_10_/k_12_/k_21_ parameters predicted by particle molecular weight (main paper, Eqs. 5-7). In each simulation, each parameter for antivenom and neutralised venom was scaled by a different randomly selected number.

We conducted bootstrap PAWN GSA on the outputs of the 2-fold simulations (Fig. S16). Application of 2-fold variability does not significantly impact the PAWN parameter rankings: the rankings remain relatively stable across all timepoints, and k_on_ is again the most influential parameter for both snakes. The GSA for the elapid envenomation case is almost identical to the simulations without the 2-fold variation. There is more visible difference with the viper envenomation case: namely that k_off_ and dose sensitivity indices overlap less, with k_off_ more influential than dose.

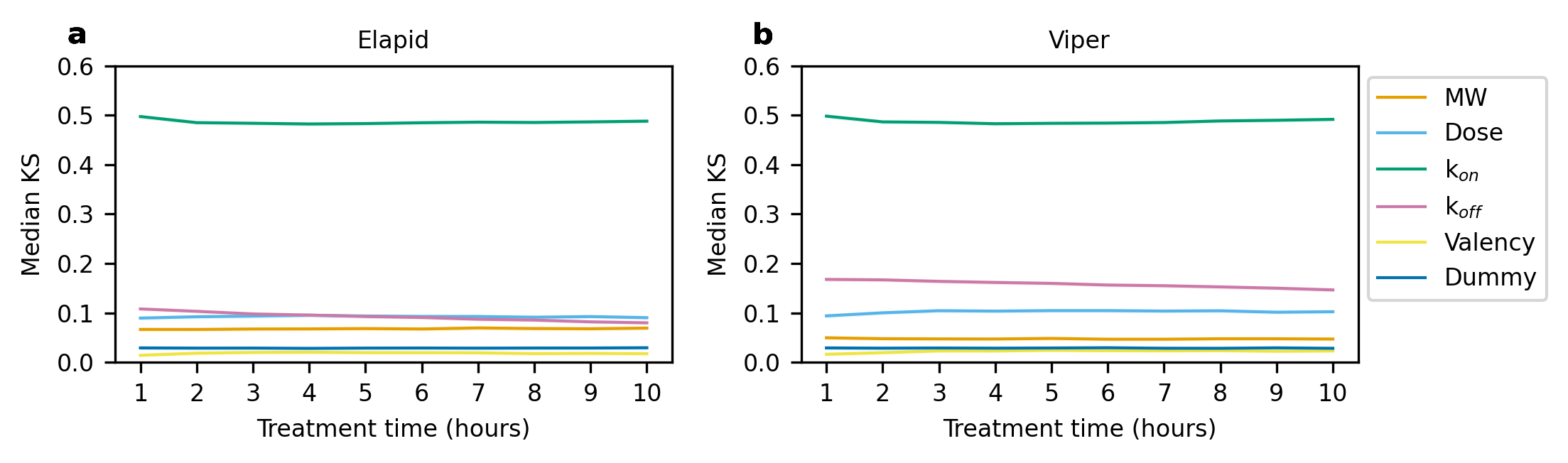

**Figure S16. PAWN GSA with two-fold variability applied to pharmacokinetic parameter prediction.**

95% Confidence intervals were plotted as clouds around the datapoint lines.

The antivenom scaffolds resulting in the 1% lowest AUC-OT scores at 1, 3, and 6 hours are shown in Fig. S17. These antivenoms occupy largely the same parameter space as the case without 2-fold variation. The same trend of narrowing parameter bounds with delayed treatment is also evident. Taken together, uncertainty in the pharmacokinetic parameter prediction does not seem to have a large impact on treatment outcome in these simulations.

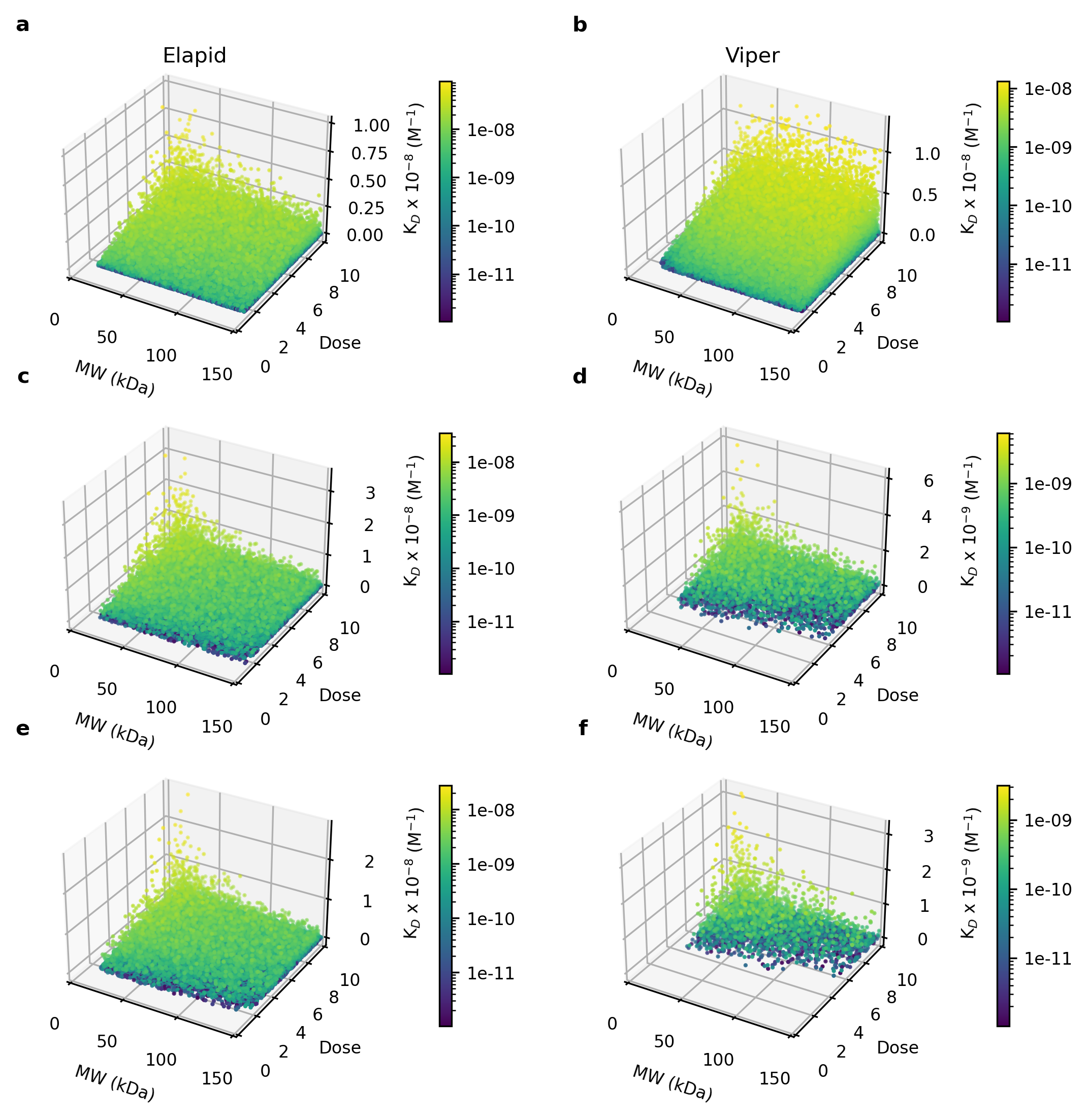

**Figure S17. Antivenom scaffolds with 2-fold variability leading to lowest 1% AUC-OT scores, at treatment times of 1, 3, and 6 hours.**

Panels on the left show elapid results. Panels on the right show viper results. Panels a-b show the results of treatment at 1 hour, panels c-d show the results of treatment at 3 hours, panels e-f show the results of treatment at 6 hours. The colour bar corresponds to antivenom K_D_ (K_D_ = k_off_/k_on_).

### 11 RBD-FAST Sensitivity Analysis

To verify the parameter ranking from the PAWN analysis, we additionally conducted a Random Balanced Design - Fourier Amplitude Sensitivity Test (RBD-FAST) on the dataset, with EASI data pre-processing. This method returns the first-order sensitivity indices (S1) of each parameter, indicating their independent effects on the model output (Tarantola et al., 2006). RBD-FAST is a variance-based method, meaning that it assumes output variation to be a valid measure output uncertainty. This assumption does not hold true for highly-skewed output distributions such as the AUC-OT output, and thus the results of this may be less reliable. EASI RBD-FAST was however chosen for validation due to its quick computation time and its tolerance of input parameters from generic sampling schemes (Goffart and Woloszyn, 2021; Plischke, 2010). EASI RBD-FAST was implemented in SALib.

The resulting RBD-FAST analysis shows largely similar trends compared to the PAWN analysis (Figure S18). While the index values for the PAWN and RBD-FAST are inherently different metrics and thus cannot be directly compared, the ranking of parameters are comparable. On the complete results distribution (Fig. S18a-b), RBD-FAST calculates k_on_ to be the most influential parameter for both envenomation cases, in agreement with PAWN. While PAWN ranks dose, k_off_, and molecular weight to have low and frequently overlapping rankings, RBD-FAST more clearly distinguishes these three parameters. Within the RBD-FAST analysis, k_off_ is the second most influential parameter, followed by dose. Molecular weight has a very small, essentially non-influential sensitivity index. We additionally conducted an RBD-FAST analysis on the sliced output array, taking 300 ng.h/ml as the threshold boundary (Fig. S18c-f). At both the head and tail of the distribution, RBD-FAST closely mimics the parameter rankings and their evolution over treatment time. Across all analyses, valency has a negligible S1 index, in line with the PAWN analysis.

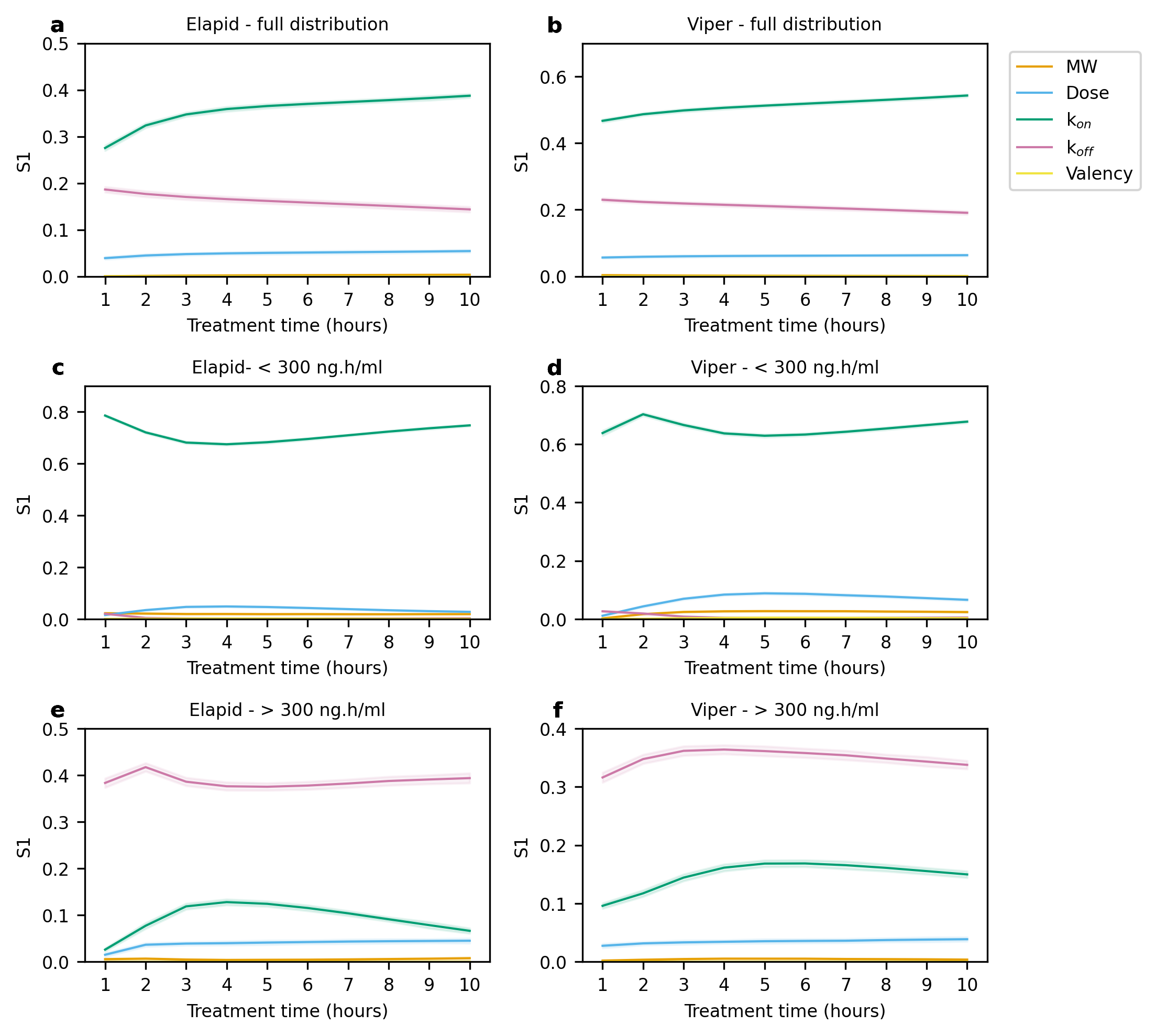

**Figure S18. RBD-FAST GSA of envenomation-treatment simulations**

S1 denotes the first-order sensitivity index. 95% confidence intervals are shown in the shaded regions. Panels on the left show elapid envenomation, panels on the right show viper envenomation. Panels a-b show the results of RBD-FAST on the complete output distribution dataset. Panels c-d show the GSA results on simulation outputs < 300 ng.h/ml, at the head of the distribution. Panels e-f show the GSA results on simulation outputs > 300 ng.h/ml, at the tail of the distribution.

### 12 k_on_ mode estimation

Within the theoretical antivenom parameter set, k_on_ sampling was performed at discrete intervals. In order to obtain a singular k_on_ mode for highly effective antivenoms, we generated KDE plots of the k_on_ distribution of the top 1% performing antivenom scaffolds. The peak of the KDE curve taken to deduce an approximate k_on_ mode. The individual KDE plots for each snake venom at each antivenom administration time point are shown across figures S19-20. In these plots, the bandwidth parameter was adjusted to ensure appropriate curve-smoothing. The KDE curves show good alignment with the approximate shape of the underlaying density histograms. With later treatment times, the k_on_ mode increases for both snake species. This is particularly noticeable for the viper envenomation case: treatment at 1 hour results in a right-skewed histogram, however increasing treatment delays shifts the k_on_ values to a left-skewed distribution.

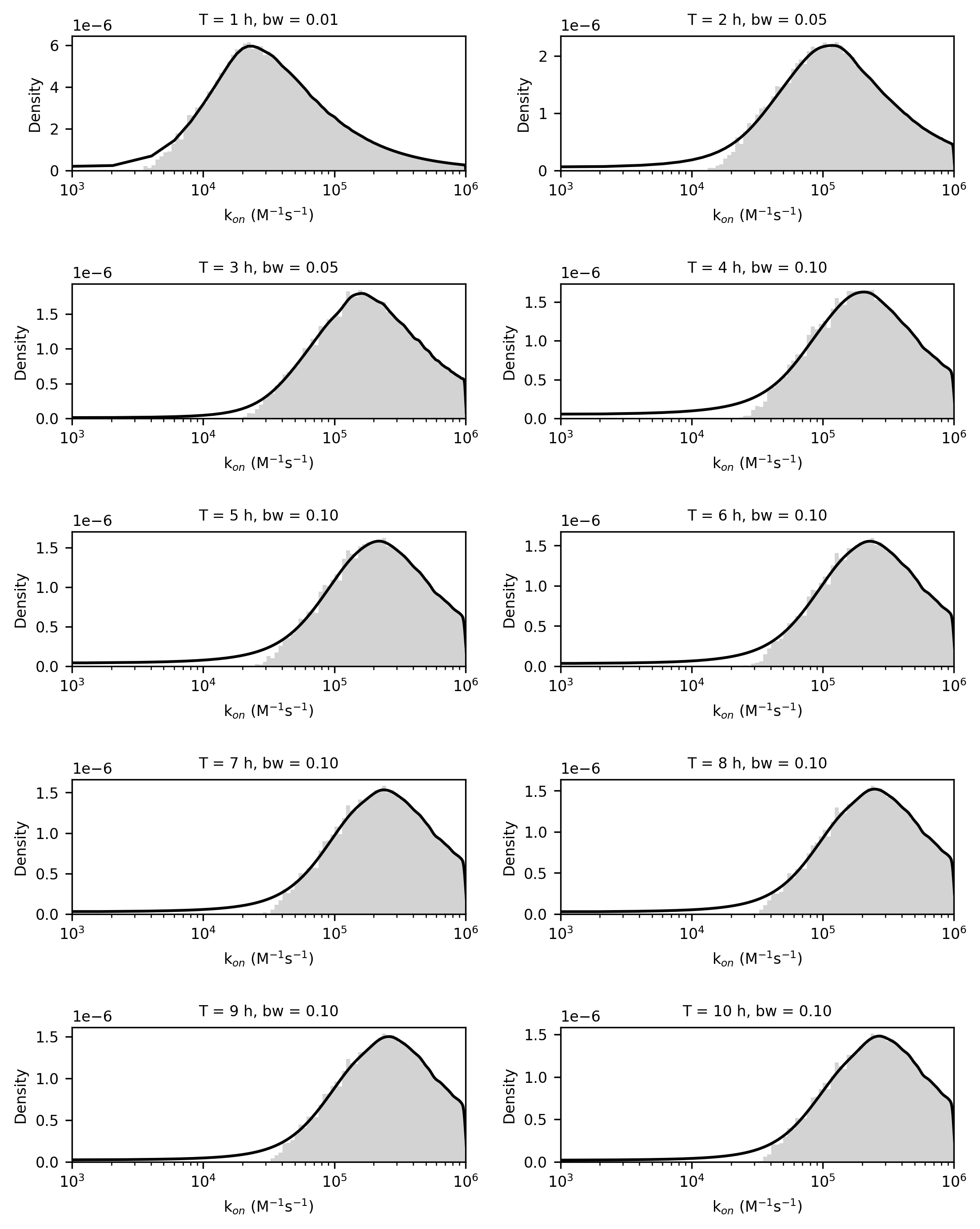

**Figure S19. Density histogram and KDE plot of k_on_ values of top 1% performing antivenom scaffolds, for elapid envenomation**

100 histogram bins taken in log-space. T = antivenom administration time (hours). bw = bandwidth.

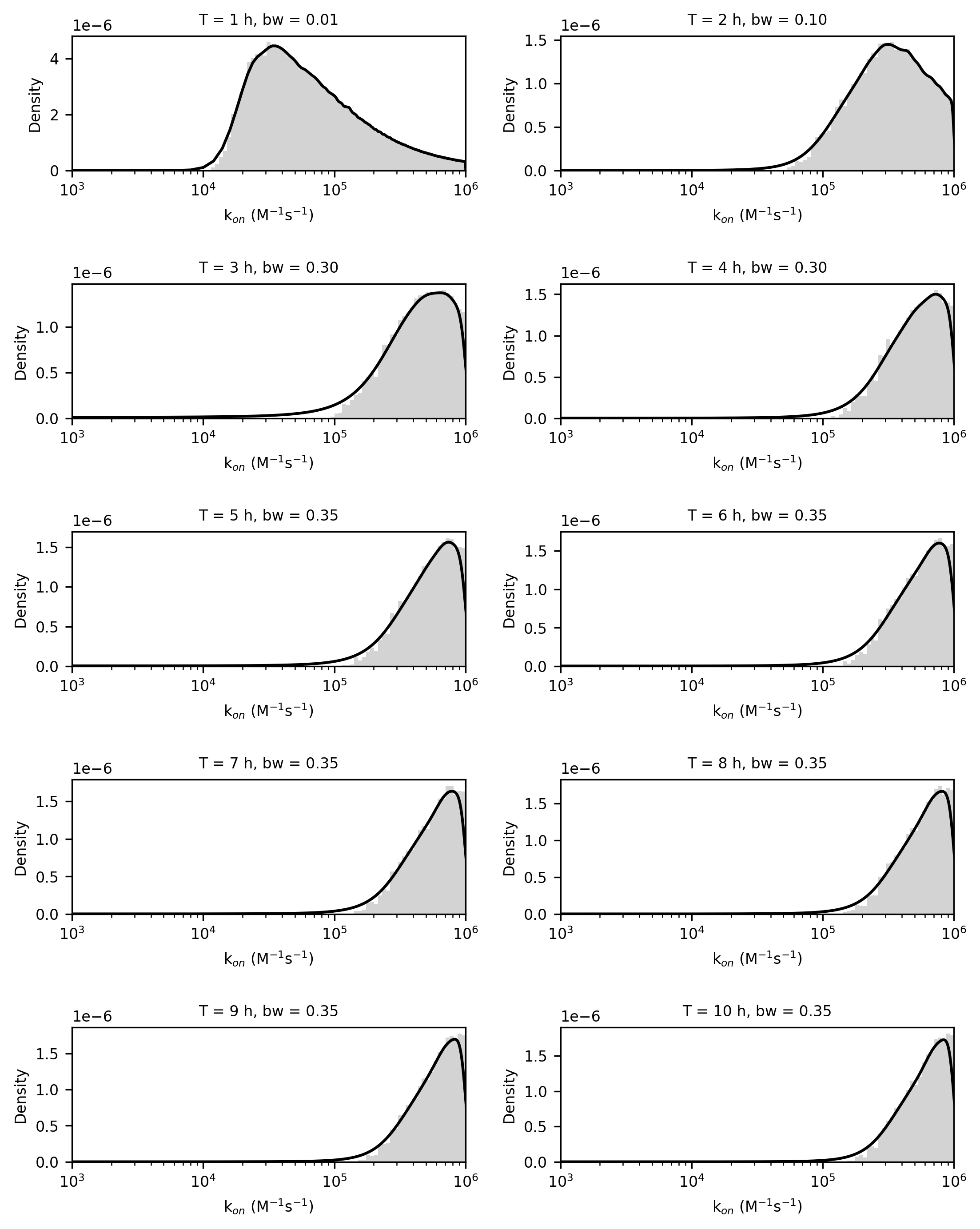

**Figure S20. Density histogram and KDE plot of k_on_ values of top 1% performing antivenom scaffolds, for viper envenomation**

100 histogram bins taken in log-space. T = antivenom administration time (hours). bw = bandwidth.

### 13 Parameter scatter plots

We plotted the sampled theoretical antivenom parameters against their resulting AUC-OT scores in the 3 hour treatment time simulations. The resulting scatter plots can indicate the strength of effect that a parameter has on the model output (Fig. S21). These plots clearly show molecular weight to have essentially no impact on model output, with an even distribution of AUC-OT scores across the entire molecular weight range. There seems to be some effect of dose on the elapid envenomation, but not on the viper envenomation. k_on_ has the largest effect for both model venoms, with high k_on_ values essential to achieve the lowest AUC-OT scores. k_off_ can be seen to have a stronger effect for the elapid venom than the viper. These scatterplots show monovalent and bivalent antivenoms taken together: separating the scaffolds by valency has a negligible impact the shapes of these plots (data not shown).

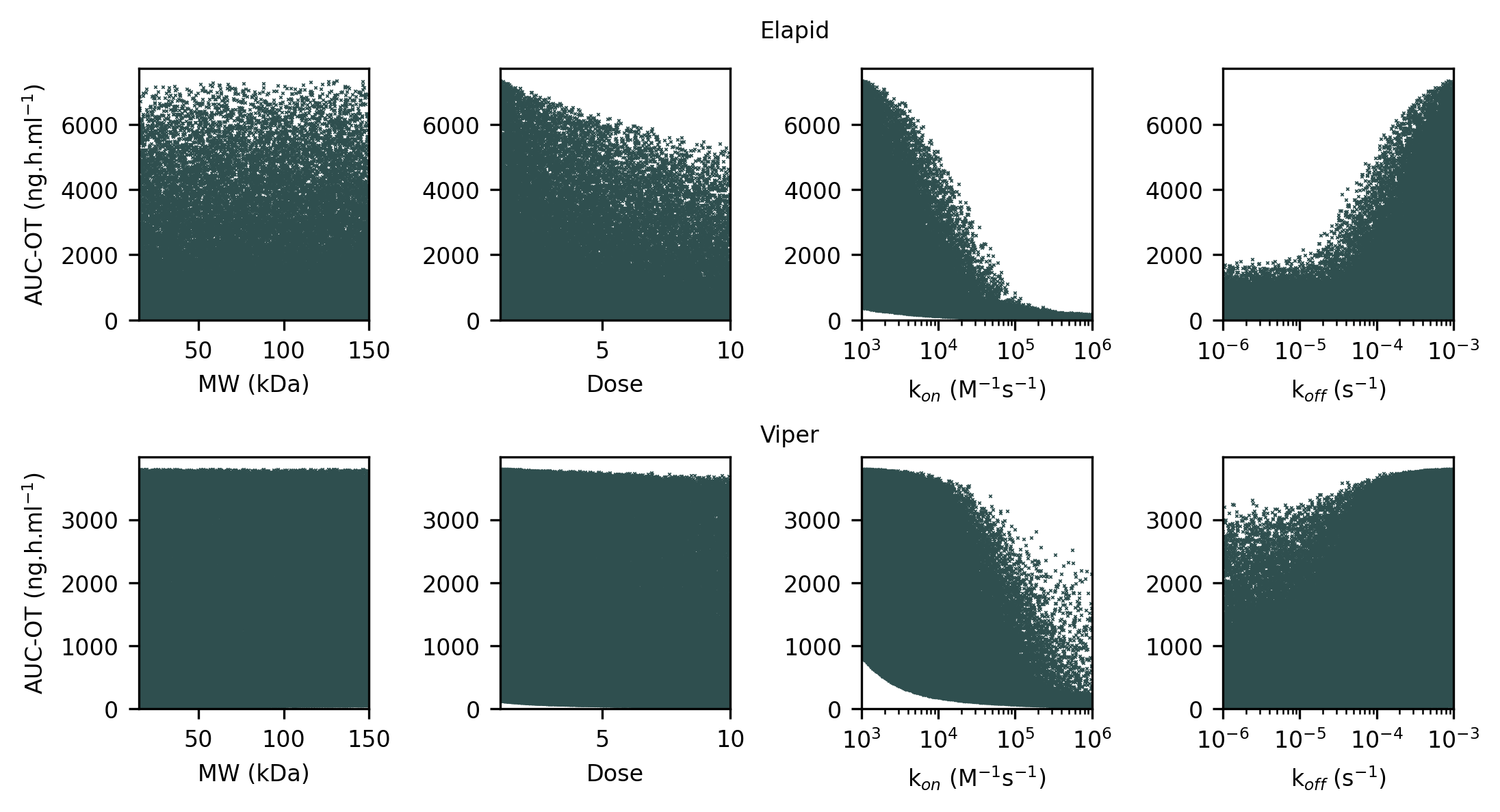

**Figure S21. Scatter plot of sampled antivenom parameters against AUC-OT outputs**

Top row shows elapid simulation results, bottom row shows viper simulation results. Data taken from 3 hour treatment time simulations.

Using the same 3 hour dataset, we next plotted the parameters pairwise, assigning colour to the resulting AUC-OT score of the scaffold. In this way, we can begin to identify parameter interactions through the presence of hot-spots that are dependent on the value of both parameters. These plots are shown in Figs. S22-23. These plots suggest an interaction between k_on_ and k_off_, since the highest values for AUC-OT essentially solely occur following treatment with antivenom scaffolds that have large k_off_ and low k_on_ values. There also appears to be a slight interaction between dose and k_on_, and dose and k_off_. In their respective pairwise plots, high AUC-OT values appear to be slightly more prevalent at combinations of low dose with low k_on_ or high k_off_.

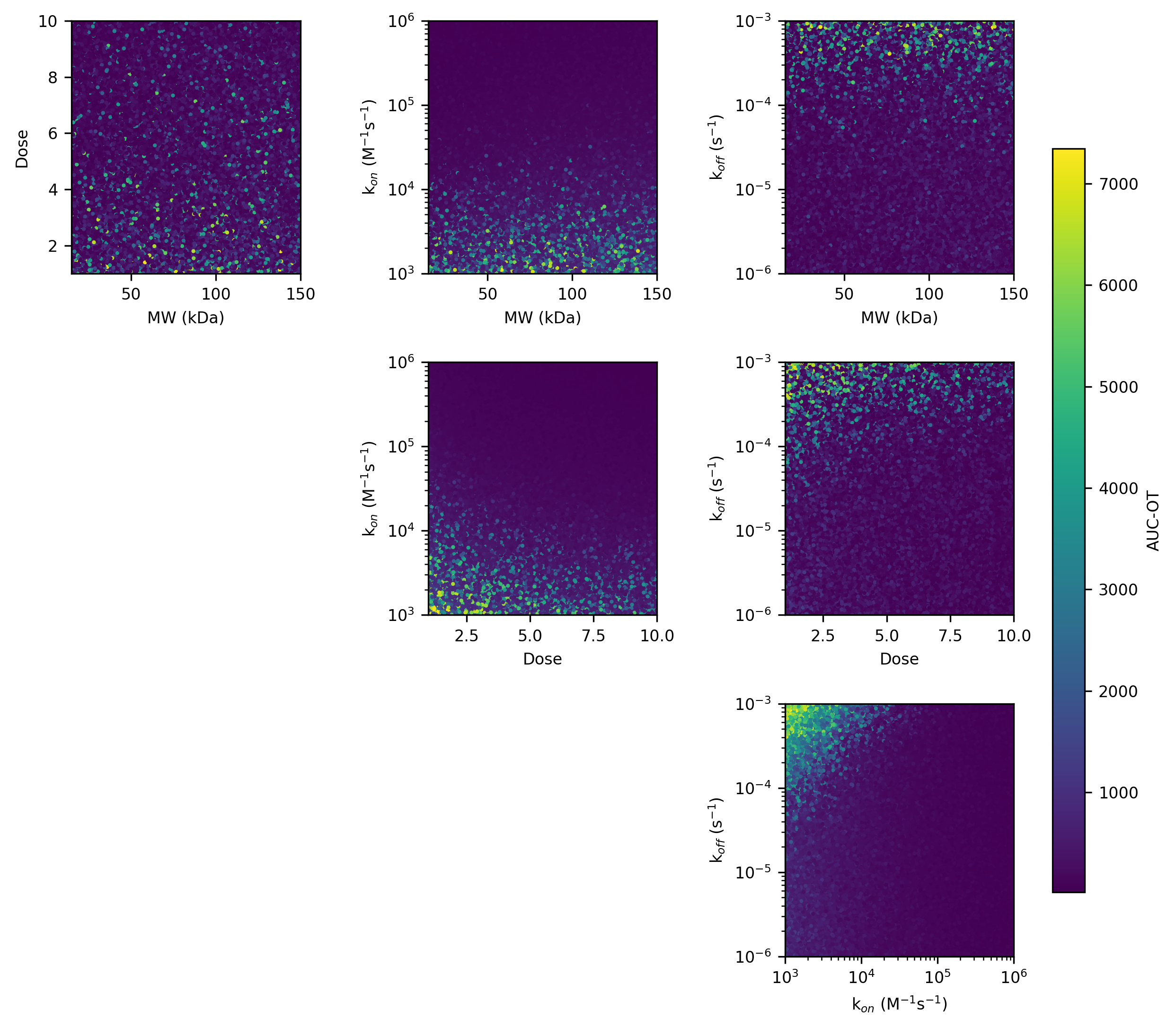

**Figure S22. Pairwise scatter plot of sampled antivenom parameters, with colour bar indicating the model elapid AUC-OT score**

Data taken from 3 hour treatment time simulations.

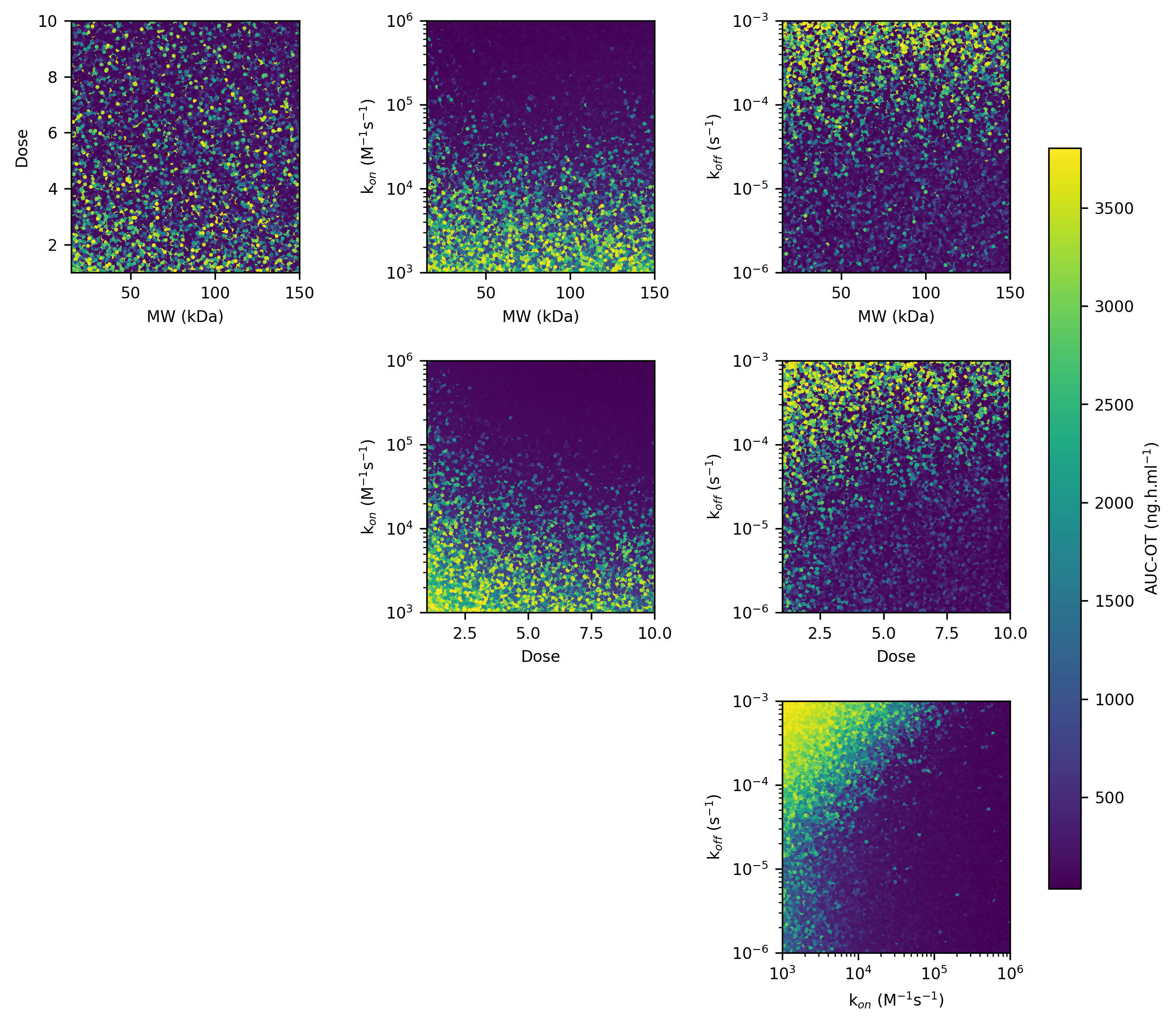

**Figure S23. Pairwise scatter plot of sampled antivenom parameters, with colour bar indicating the model viper AUC-OT score**

Data taken from 3 hour treatment time simulations.
